# Autophagy maintains HEV identity and function during inflammation

**DOI:** 10.1101/2025.08.06.668862

**Authors:** Kathryn A Jacobs, Yichao Hua, Apostolis Panagiotis Nikolakopoulos, Swati Gupta, Gerlanda Vella, Ana Filipa Domingues Laranjeiro, Julie Bonnereau, Mélanie Guyot, Larissa Mourao, Feyza Cansiz, Gabriele Allies, Nicolas Van Baren, Sophie Lucas, Wiebke Sondermann, Florian Rambow, Alpaslan Tasdogan, Colinda Scheele, Patrizia Agostinis, Gabriele Bergers

## Abstract

High endothelial venules (HEVs) play a crucial role in adaptive immune responses in secondary and tertiary lymphoid organs. They are uniquely equipped with high levels of peripheral node addressins (PNAd), harboring carbohydrate structures that serve as L-Selectin ligands to efficiently facilitate lymphocyte homing. During inflammation, the HEV network expands in SLOs, increasing lymphocyte infiltration, but the underlying mechanisms maintaining HEVs remain underexplored. Here, we report that autophagy is essential for HEV function and expansion. Using single-cell transcriptomics, intravital imaging, and an inducible HEV tracer system in mice, we demonstrate that autophagy deficiency compromises LTβR-signaling and the Unfolded Protein Response in HEVs, leading to disrupted PNAd production, dedifferentiation, and reduced lymphocyte homing. Autophagy deficiency and LTβR blockade impaired HEV function and reduced skin inflammation in psoriasis-bearing mice by limiting immune infiltration and cytokine release. Our work uncovers an unprecedented role of autophagy in safeguarding HEV identity and function during inflammation.

**Highlights:**

- High endothelial venules (HEVs) exhibit heightened autophagy in comparison to non-HEV blood endothelial cells, which further increases during inflammation.
- Autophagy is pivotal in maintaining HEV fate and function, specifically during inflammation, via LTβR signaling and the Unfolded Protein Response (UPR), ensuring the proper production of peripheral node addressins (PNAd) that serve as L-selectin ligands for the efficient influx of naïve lymphocytes.
- Blocking autophagy in HEVs leads to disrupted PNAd production, HEV flattening and dedifferentiation, and reduced lymphocyte homing.
- Genetic and pharmacological perturbation of HEV function in the lymph nodes and skin lesions of psoriasis-bearing mice impaired neutrophil and lymphocyte recruitment, as well as cytokine secretion, thereby alleviating skin inflammation.

## Introduction

High endothelial venules (HEVs) are specialized organotypic endothelial cells in secondary lymphoid organs, such as lymph nodes (LN), that serve to facilitate naïve lymphocyte entrance into the LN parenchyma for antigen priming. These endothelial cells have a distinctive cuboidal morphology in contrast to the flat appearance of other blood endothelial cells (BECs) and express uniquely decorated sialomucins. These L-selectin ligands carry high levels of peripheral node addressin (PNAd), glycoproteins with complex carbohydrate structures containing 6-sulfo sialyl Lewis^X^-capped O-glycans that bind L-selectin on naïve lymphocytes, efficiently facilitating their passage through HEVs^1–3^. In response to infections and inflammation, HEVs expand their PNAd+ network and upregulate additional leukocyte adhesion molecules such as P-selectin, E-selectin, Icam1, and Vcam1 to increase lymphocyte influx for priming^4^. HEVs can also be ectopically induced in non-lymphoid tissue at sites of inflammation during infection, autoimmunity, and graft rejection, as well as in cancer, where they support the formation of tertiary lymphoid structures ^5–10^. While HEVs in cancer are associated with a better immune response and subsequent prognosis^1,8^, they exacerbate the phenotype and symptoms during chronic inflammatory diseases. The key signaling pathway involved in HEV formation is the lymphotoxin beta receptor (LTβR) and downstream non-canonical NF-κB axis, which is activated by Lymphotoxin α1β2 (LTα1β2) or LIGHT (TNFSF14), produced by lymphocytes and dendritic cells. This induces the expression of genes involved in lymphoid organogenesis, such as those encoding HEV-specific enzymes for PNAd construction and chemokines that attract lymphocytes^11^.

Macroautophagy (hereafter referred to as autophagy) is a major lysosomal degradation pathway with an essential role in cellular homeostasis. During autophagy, cellular components, either in bulk or selectively, are enclosed within autophagosomes and trafficked to the lysosome, where, upon fusion, the cargoes are broken down and recycled to fulfill the cell’s metabolic needs^12^. While most cells maintain a basal level of autophagy, during periods of cellular or environmental stress, they upregulate this pathway to clear damaged organelles and protein aggregates, as well as to provide necessary metabolites^12^. Overall, autophagy is considered an anti-inflammatory process, as it can directly degrade pathogens and other inflammatory mediators. Moreover, loss of autophagy in most inflammatory conditions increases proinflammatory phenotypes^13^. In inflamed cremaster muscle BECs, a lack of autophagy leads to increased expression of adhesion molecules and increased neutrophil diapedesis across venules^14^. A similar mechanism is at play in BECs of melanoma-bearing mice, where lack of autophagy leads to increased Icam1 and Vcam1 expression on tumor vessels, and subsequent intratumoral infiltration of cytotoxic CD8+ T-cells, which fostered anti-tumor immunity^15^.

Complementing autophagy, during prolonged cellular stress, the Unfolded Protein Response (UPR) tackles the accumulation of misfolded proteins in the endoplasmic reticulum (ER) via ER stress sensors Perk, Ire1α, and Atf6, which results in increased transcription of genes involved in protein folding and degradation^16^. Intriguingly, all three ER stress sensor pathways promote autophagy by increasing the expression of autophagy-related genes^16,17^. The Ire1α pathway specifically promotes autophagy in endothelial cells, through its downstream mediator, the transcription factor XBP1s, which binds to the promoter of the essential autophagy gene *Beclin1* and induces its transcription. While the UPR has been extensively shown to activate autophagy, ER stress can also negatively affect autophagy in pathological contexts. In models of neurodegenerative diseases, knock-out of UPR components resulted in increased autophagy activation, suggesting the UPR is dampening autophagy signaling^18^. There is thus a clear interlinkage between both pathways.

Our previous transcriptomics analysis of endothelial cells (EC) and HEVs in tumors, as well as HEVs in homeostatic lymph nodes (LN), revealed heightened autophagy-related pathways in HEVs ^8^. Autophagy can regulate lipid dynamics and membrane organization, providing metabolites essential for maintaining membrane structure^19^. Since a key feature of HEVs is PNAd production, we posited that autophagy may influence the composition of these surface glycan molecules.

Here, we uncover an exceptional role for autophagy in maintaining HEV function and identity. Employing unique HEV tracer models, intravital imaging, and single-cell transcriptomics, we demonstrate that genetic ablation of the essential autophagy genes ATG12 or ATG7 in HEVs was associated with reduced expression of UPR pathway genes and LTβR activity, leading to HEV dedifferentiation and reduced lymphocyte influx in LNs and psoriasis skin lesions. Thus, targeting HEVs can attenuate both systemic and local adaptive immune responses in inflamed tissues.

## Results

### Autophagy maintains HEV function in response to acute, inflammatory stress

To gain insight into autophagy-dependent effects on HEV fate and function during inflammation, we utilized the well-established oxazolone (OX) skin painting model, which creates an acute contact dermatitis and LN reaction in mice with a peak of inflammation on day 3 post-skin painting, and a resolution by day 7 (**Figure 1A**)^4,20^. Single-cell RNA sequencing (scRNAseq) analysis of homeostatic LN HEVs versus inflamed OX-D3-treated HEVs revealed increased autophagy activity in inflamed HEVs (**Figure 1A, S1A**). We confirmed autophagosome occurrence in homeostatic HEVs by detection of Lc3b puncta (autophagosomes) on LN sections, which increased in HEVs of OX-D3-treated LNs, and decreased to baseline levels at OX-D7 when inflammation is resolving (**Figure S1B, C**). We further detected an increased autophagic flux in inflamed LN HEVs in comparison to homeostatic HEVs by employing the Lc3b GFP-RFP reporter mouse in which GFP is quenched when autophagosomes are degraded in the lysosome^4,21,22^, therefore giving a lower ratio of GFP to RFP when autophagy is more active (**Figure 1B, C**). Notably, this phenotype was HEV-specific because not only did HEVs exhibit higher levels of autophagosomes, but also they increased autophagy in LNs during inflammation, whereas non-HEV BECs did not, as quantified by Lc3b puncta and transcriptomics (**Figure 1D-H).** Similar results were obtained from human tonsils, which are HEV+ SLOs. Using a publicly available scRNAseq dataset of stromal cells in human tonsils, we identified 6 subclusters of BECs, of which one comprised HEVs. We identified 81 HEVs that expressed 6 of the 7 genes of our in-house and published human HEV signature (CHST4, FUT7, GCNT1, CD24, CH25H, ENPP6 B3GNT3 and TMEM176A)^8^(**Figure S1D-G**). Using published autophagy activation and macroautophagy gene signatures ^23^, we found that HEVs in human tonsilitis displayed a trend towards increased autophagy activation signature compared to homeostatic HEVs (**Figure S1H**) and that autophagy was significantly increased in HEVs versus BECs (**Figure 1I-K**). In line, tonsilitis biopsies from 8 patients revealed more Lc3b puncta in HEVs than in non-HEV BECs (**Figure1L, M**). Together, these data infer heightened autophagy activity in HEVs compared to non-HEV BECs, which is further elevated during inflammation in murine and human SLOs.

**Figure 1:**
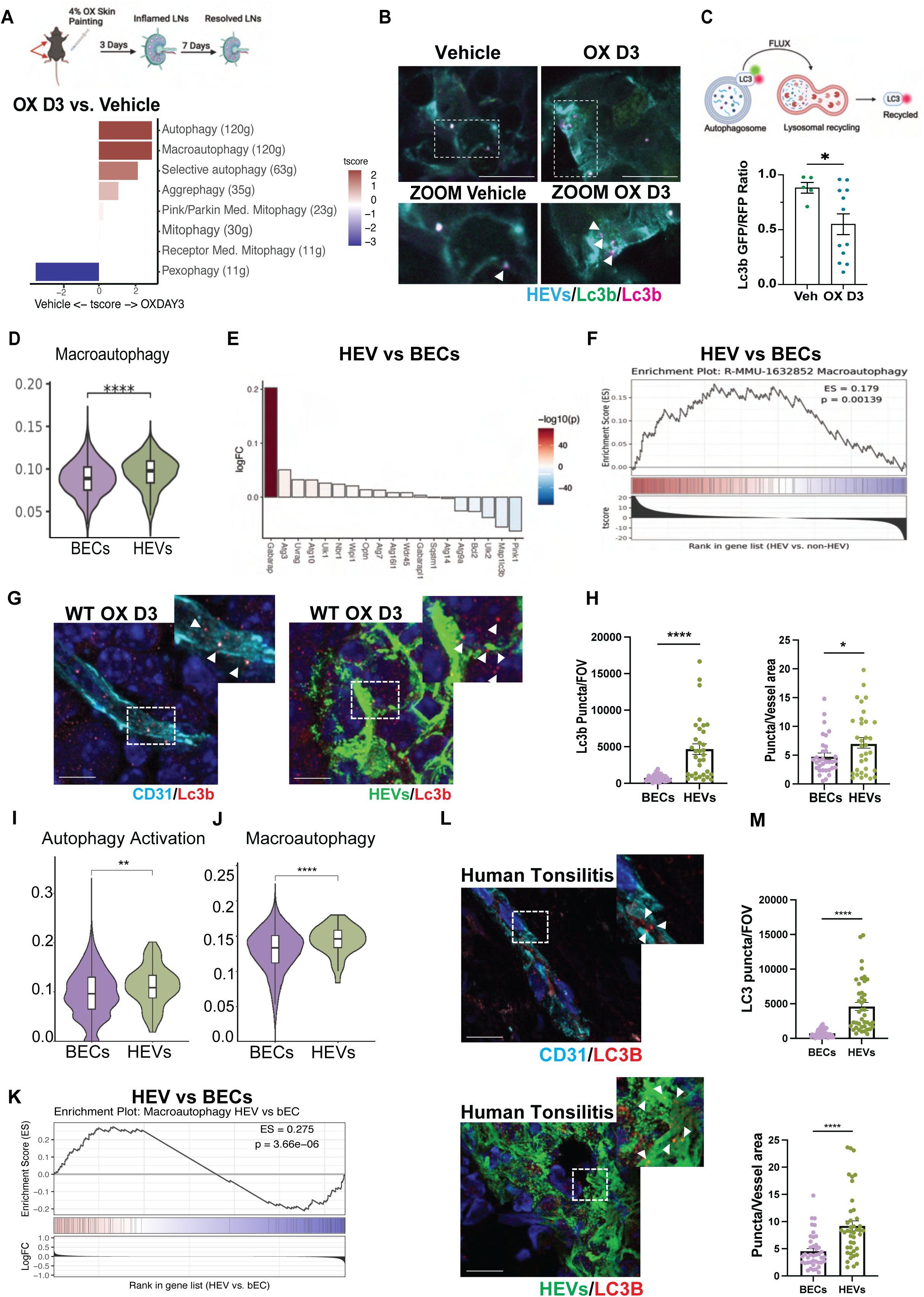
Autophagy is upregulated in HEVs compared to BECs and further upregulated during inflammation. (A) Treatment schedule of OX (top) and Waterfall plot of different Reactome autophagy-related signatures comparing steady state (vehicle) and OX-treated WT HEVs (bottom) (B) Immunofluorescence staining of Meca-79 (HEVs) and RFP-Lc3, GFP-Lc3 reporter in Vehicle or OX treated mice. Scalebar= 10um Arrows denote single positive RFP-puncta. (C) Scheme and quantification of the ratio of GFP/RFP double-positive puncta to RFP single positive puncta representing autophagic flux. (D) Violin plot of Reactome macroautophagy signature in BECs vs HEVs from vehicle and OX-treated mice (E) Waterfall plot of genes associated with autophagy activation in HEVs vs non-HEV ECs from vehicle and OX-treated mice (F) GSEA plot for macroautophagy in HEVs vs BECS from vehicle and OX-treated mice (G) Immunofluorescence staining of Lc3b in inflamed (OX D3) HEVs or BECs (CD31), Scale bar=10um (H) Quantification of (G) (I) Violin plot of active autophagy signature in BECs vs HEVs from homeostatic and inflamed human tonsils (J) Violin plot of macroautophagy signature in BECs Vs HEVs from human tonsils from homeostatic and inflamed human tonsils (K) GSEA plot of macroautophagy signature in HEVs vs. BECs from homeostatic and inflamed human tonsils (L) Immunofluorescence staining of BECs (CD31), HEVs, and LC3B in human tonsilitis tissue. Scalebar =10um (M) Quantification of LC3B puncta in (L) *p < 0.05; **p < 0.01; ***p < 0.001; ****p < 0.0001.

To assess the functional significance of autophagy in LN HEVs, we conditionally knocked out the essential autophagy genes *Atg12* or *Atg7* in the EC compartment by crossing ATG12 or ATG7 fl/fl mice with Cdh5-CreERT2 mice expressing a tamoxifen-inducible Cre recombinase (ATG12^ECKO;^ ; ATG7^ECKO^). We first addressed the extent to which autophagy-impaired HEVs were still able to respond to inflammation. As anticipated, the common endothelial adhesion molecules P-selectin, Icam1, 4-1BB, as well as vWF, were elevated to differing degrees in OX-D3-HEVs (**Figure 2A, B**). The inflammatory response, however, was compromised in HEVs of autophagy ATG12^ECKO^ and ATG7^ECKO^ mice as reflected in the diminished upregulation of these inflammation molecules (**Figure 2A, B, S2A-D**).

**Figure 2:**
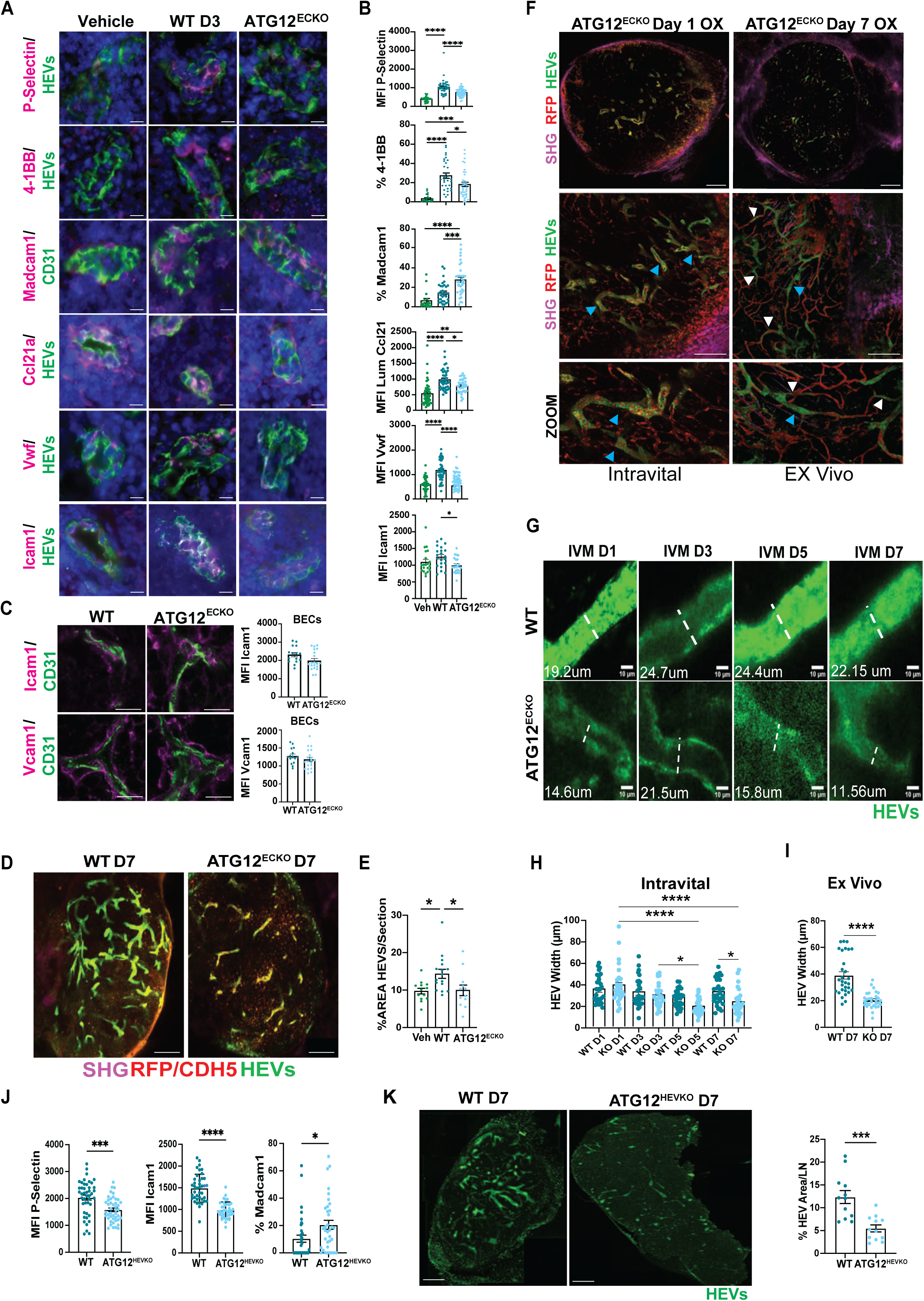
autophagy is essential to HEV maintenance in response to inflammatory stress. (A) Immunofluorescence staining of HEVs (Meca-79), Cd31, P-selectin, 4-1BB, Madcam1, Ccl21a, Vwf, and Icam1 in Vehicle or OX D3 treated WT and ATG12^ECKO^ mice, scalebar=10um (B) Quantification of stainings in (A) (C) Immunofluorescence staining and MFI quantification of CD31 and Icam1 or Vcam1 in BECs on Day 3 post-OX treatment. Scale bar =10um (D) Immunofluorescence imaging of HEVs and CD31 (WT) or endogenous RFP (ATG12^ECKO^) on Day 7 post-OX treatment scale bar =250um (E) Quantification of Meca-79 (HEV) Area per LN section in WT vs ATG12^ECKO^ mice on OX D7 (F) Representative immunofluorescence imaging of HEVs, endogenous RFP, and collagen (SHG) in an ATG12^ECKO^ mouse on Day 1 post OX treatment (intravital) and the same mouse on Day 7 post-OX treatment (Ex Vivo) scale bar =200 um. Blue arrows show mature HEVs; white arrows denote flattened HEVs which lose PNAd. (G) Representative HEVs (Meca-79, intravital) over time (Day 1-7 post-OX treatment) and corresponding vessel width in a WT and ATG12^ECKO^ mouse. Scale bar =10um (H) Quantification of the width of 20 representative HEVs over time (Day 1-7 post-OX treatment) in WT and ATG12^ECKO^ mice (I) Quantification of HEV width on Day 7 post OX treatment Ex vivo in WT and ATG12^ECKO^ mice. (J) Quantification of MFI P-selectin, MFI Icam1 and percentage of Madcam1-positive HEVs in WT vs ATG12^HEVKO^ mice on OX D3. (K) Immunofluorescence imaging of HEVs and quantification of HEV area per whole LN in WT vs ATG12^HEVKO^ mice on OX D7. Scale bar = 250um *p < 0.05; **p < 0.01; ***p < 0.001; ****p < 0.0001.

Surprisingly, this response in LNs was HEV-specific, as Icam1 and Vcam1 were equally expressed in inflamed autophagy-proficient or deficient non-HEV BECs in LNs (**Figure 2C**). Furthermore, the chemokine Ccl21a, which binds to the luminal surface of activated HEVs to attract more lymphocytes during inflammation^4^, predominantly remained on the abluminal side, and overall levels were reduced in autophagy-compromised HEVs (**Figure 2A, B, S2A, B**). These results point to a context and endothelial subtype-specific role of autophagy in HEVs^15^. Interestingly, the impaired HEV adaptation response to inflammation in ATG12^ECKO^ and ATG7^ECKO^ LNs correlated with a significant increase of the integrin ligand Madcam1, known to be transiently expressed during peripheral LN development in HEVs and in immature HEVs of peripheral LN^24^ (**Figure 2A, B, S2A, B**). These results suggest that autophagy-impaired HEVs may become immature and potentially undergo dedifferentiation, which should be reminiscent of a flattened endothelial and thinned vessel appearance, concomitant with the loss of PNAd.

Thus, we evaluated the LN HEV architecture on day 7 post-OX -treatment in wildtype (WT) and knockout (KO) mice. While the HEV network in wildtype LN had expanded in response to OX to accommodate increased lymphocyte infiltration, autophagy deficiency in HEVs prevented an increase in their network (**Figure 2D-F, S2E-F**). Notably, HEVs lost their characteristic architecture and appeared thinner and less plump (**Figure 2D, F, G, S2E**). In addition, we followed HEV expansion over time in WT and ATG12^ECKO^ mice by intravital microscopy using an imaging window over the inguinal LN (**Figure 2F-I**). In contrast to wildtype HEVs, following ATG12^ECKO^, HEVs in LNs from day 1 to 7 revealed a flattening HEV appearance with a decreasing vessel width and a lack of HEV expansion over time in the HEV compartment (**Figure 2F-I).** Of note, on day 7 post-OX, Madcam1 levels in HEVs remained elevated in ATG12^ECKO^ mice, and Ccl21 levels remained low and abluminal (**Figure S2F**). To further strengthen the HEV-specific effects, we genetically ablated ATG 12 only in HEVs, using a Chst4-CreER reporter mouse. This reporter enables the ATG12 gene knockout in approximately 40-60% of HEVs, in concordance with the distribution and levels of Chst4 in HEVs. Even in this chimeric situation, we observed a similar impaired response during inflammation as shown by the reduced P-Selectin and Icam1 and increased Madcam1 levels in ATG12KO HEVs (**Figure 2J**). This phenotype is also opposite to the effects we previously observed in tumor ECs, in which autophagy deficiency increased inflammatory adhesion molecules such as Icam1 and Vcam1^25^. Moreover, these mice displayed similar thinning on day 7 compared with the full endothelial knockout (**figure 2K**). Together, these data demonstrate that autophagy-deficient HEVs lose their phenotypical characteristics and lose their responsiveness to inflammation.

### Autophagy deficiency causes HEV reprogramming and transdifferentiation

To gain more mechanistic insight into the autophagy-regulated HEV identity in response to inflammation, we isolated CD45-CD31+ ECs from peripheral LNs of WT and ATG12^ECKO^ mice, treated with vehicle or OX for 3 and 7 days, and performed 10x Genomics single-cell transcriptomics. We identified 11 subclusters from the obtained 22272 ECs reflective of lymphatic EC (2), capillary ECs(4), arterial EC (1), venous EC (1), HEVs (1), and mitotic EC (1) as further validated by differentially expressed gene (DEG) analysis (**Figure 3A, S3A**). We then employed SCENIC to identify the top transcription factors (TF) with the highest activity in HEVs compared to other LN ECs (**Figure 3B, S3B**). In line with the previously identified TFs in LN HEV and ectopic TU-HEVs^4,8,26,27^ (**Figure S3B**), we identified RelB and NFκb2, the key transcription factors in the LTβR/non-canonical NFkB axis. Interestingly, besides Hox, Jun, and Ets TF family members, we also noted high x-box binding protein-1 (Xbp1) activity, which is downstream of UPR sensor IRE1a system in response to ER stress^28^. Strikingly, most of the top TF activities were downregulated in autophagy-compromised HEVs on OX-D3, and all were suppressed on OX-D7, suggestive of significant reprogramming in HEVs during inflammation in the absence of autophagy (**Figure 3B**). Based on our observation that PNAd levels were also reduced in ATG12^ECKO^ HEVs (**Figure 2F**), we investigated the expression level of adhesion molecules and the essential enzymes that are involved in the production of PNAd in LN HEVs of WT and ATG12^ECKO^ mice on OX-D3 and OX-D7 (**Figure 3C**). We noticed a marked decrease in the expression of *Glycam1*, *Chst4, Chst2*, *Gcnt1, St3gal4, St3gal6*, and Fut7 on OX-D3, with a continuing decrease of *Chst4, Chst2, Fut7, Gcnt1 and Glycam1* on OX-D7 treatment in ATG12^ECKO^ HEV (**Figure 3C, S3C**). Interestingly, the resultant autophagy-compromised HEVs resembled those of LN HEVs that were treated with an LTβR antagonist ^4^, conceivably due to the observed suppression of the downstream effectors of LTbR acting NFkB2/RelB transcription factors. Based on these results, we hypothesized that the absence of autophagy may trigger HEV reprogramming in part by inhibiting LTbR signaling.

**Figure 3:**
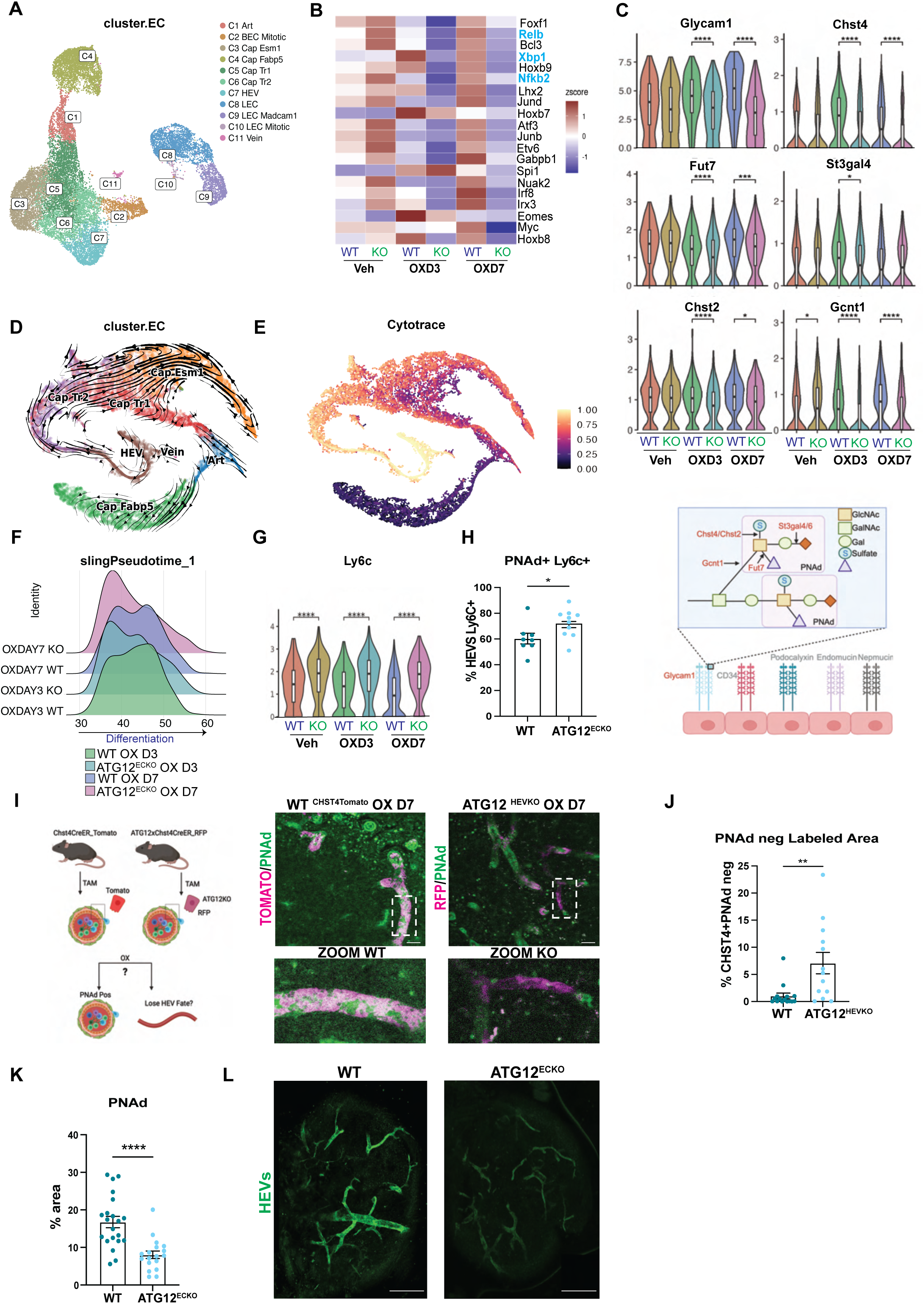
During inflammation HEVs lose their mature identity in the absence of autophagy. (A) UMAP plot colored by unsupervised EC clustering in WT and ATG12^ECKO^ vehicle, OX Day 3 and OX Day 7 mouse LN (B) TF activity of Top transcription factors in HEVs as predicted by SCENIC (C) (top) Violin Plots of selected enzymes involved in PNAd production across samples. (Bottom) Schematic representation of PNAd structure and the enzymes involved in its production. (D) Differentiation trajectory of the lymph node vasculature as predicted by Velocyto/ScVelo. Differentiation direction is indicated by arrows in the entire datasets. (E) Differentiation potential predicted by CytoTRACE on the trajectory in (D). (F) Graphical representation of slingshot analysis of HEV cluster plotted in pseudotime for WT and ATG12^ECKO^ on OX Day 3 and OX Day 7. (G) Violin plots for Ly6c in HEVs across samples (H) % of HEV cells analyzed by FACs which are double positive for PNAd (Meca79) and Ly6C. Each dot represents one mouse. (I) (Left) Schematic representation of tracer experiment. (Right)immunofluorescence staining of HEVs (PNAd) and CHST4 Tracer (Tomato for WT or RFP for ATG12^HEVKO^) in OX Day 7 treated LNs. Scalebar = 50um. (J) Quantification of (I) reporter positive cells (Tomato for WT or RFP for ATG12 ^HEVKO^) which are PNAd-negative. (K) Quantification of HEV area in Steady-state WT vs ATG12^ECKO^ mice 6 weeks post Tamoxifen treatment. (L) 3D reconstruction of immunofluorescence imaging of HEVs in Steady-state WT vs ATG12^ECKO^ mice 6 weeks post Tamoxifen treatment. Scale bar =250um *p < 0.05; **p < 0.01; ***p < 0.001; ****p < 0.0001.

Palantir, scVelo, and CytoTRACE analyses revealed a trajectory in which HEVs are positioned between Cap Tr2 EC and veins **(Figure 3D,E**). Of all EC subpopulations, HEVs have surprisingly the highest differentiation potential next to Cap TR2 ECs. Interestingly, this cluster contains transitional ECs that express canonical capillary genes such as Ly6c as well as some HEV genes, albeit at low levels compared to mature HEVs, and are a source for HEV neogenesis after immunization^26^ (**Figure 3E**). Congruently, slingshot pseudotime analysis exposed two differentiation peaks over time in OX-D3 and OX-D7 WT HEV, which may reflect differing maturation stages of HEVs. In contrast, ATG12-KO HEVs on OX-D3 and even more strikingly on OX-D7, shifted back in pseudotime, predicting a reverted phenotype towards a less differentiated cell state **(Figure 3F, S3D)**. To test whether ATG12-KO HEVs transition into capillary cell state, we investigated the expression of the capillary marker *Ly6c* in the HEV cluster of our single-cell data. In support, *Ly6c* expression was elevated in ATG12^ECKO^ HEVs, but not in WT HEVs, at OX-D3 and OX-D7. (**Figure 3G**). Further confirmation stems from the detection of an increased transitory MECA79+Ly6c+ HEV population during inflammation by Flow cytometry analysis in the LNs of ATG12^ECKO^ mice, suggesting that HEVs may differentiate into Cap TR2 ECs (**Figure 3H, S3E**).

Finally, to prove that autophagy ablation in HEVs causes loss of HEV identity over time, we traced and mapped the fate of HEVs *in vivo*. Using our generated ATG12^HEVKO^ mice, which upon tamoxifen treatment, labels the Chst4-expressing ATG12-KO HEVs with RFP (**Figure 3I**). Although we only observed about 60% recombination efficiency due to the low and varying Chst4 expression levels in HEVs, we were able to detect significant differences between WT and ATG12-KO HEVs (**Figure 3I-J**). As a WT control, we crossed the Chst4 CreER reporter mice with Rosa-CAG-LSL-tdTomato mice in which WT-HEV express tdTomato upon tamoxifen induction. We then treated both groups with OX and traced PNAd+, tdTomato+ WT-HEVs and PNAd+ RFP+ ATG12-KO HEVs to determine how many HEVs in both groups would become PNAd-negative. While we found very few PNAd-tomato+ cells in the WT condition (average 1%), we noticed up to 23% (average 7%) PNAd-negative RFP/Tomato+ cells in LNs of ATG12^HEVKO^ mice (**Figure 3I, J**). We further confirmed that these changes were not caused by apoptosis or a reduced proliferation rate (**Figures S3F, G**). It is essential to note that the ablation of autophagy in homeostatic HEVs also resulted in a loss of HEV identity, characterized by the same phenotypes of PNAd loss, and reduced HEV area with thinner HEVs (**Figure 3K,L**). It required, however, 6 weeks to observe these changes, probably due to the low turnover rate of homeostatic HEVs (**Figure 3K, L**). Taken together, these data uncover that HEVs require autophagy to maintain their identity, especially during inflammation, as they lose PNAd and transition into a flattened capillary cell state when autophagy is ablated.

### Autophagy-deficiency in LN HEVs compromises adaptive immune response

Does HEV transdifferentiation and loss of HEV fate then impact their ability to facilitate the heightened lymphocyte influx during inflammation? We performed flow cytometry analysis of the different adaptive and innate immune cell populations in LNs with autophagy-proficient or deficient HEVs at OX-D3 and OX-D7 compared to homeostasis (vehicle). As anticipated, during the inflammation peak at OX-D3, the overall CD45+ cell population, and the subpopulations of naïve and activated CD8 and CD4 T-cells, Tregs, as well as B-cells and dendritic cells (DC) were significantly increased in autophagy-proficient HEV conditions (**Figure 4A-H, S4A, B**). In contrast, LNs harboring autophagy-deficient HEVs showed a dampened influx of lymphocytes compared to the WT condition (**Figure 4A-H).** Similar trends were seen with B-cells, whereas the number of dendritic cells (DC) remained unaffected in LNs of ATG12^ECKO^ mice, as DCs predominantly enter the LN via lymphatics and not via HEVs (**Figure S4A, B**). At OX-D7, when inflammation ceased, lymphocyte influx decreased and were similar in LNs of WT and ATG12^ECKO^ mice.

**Figure 4:**
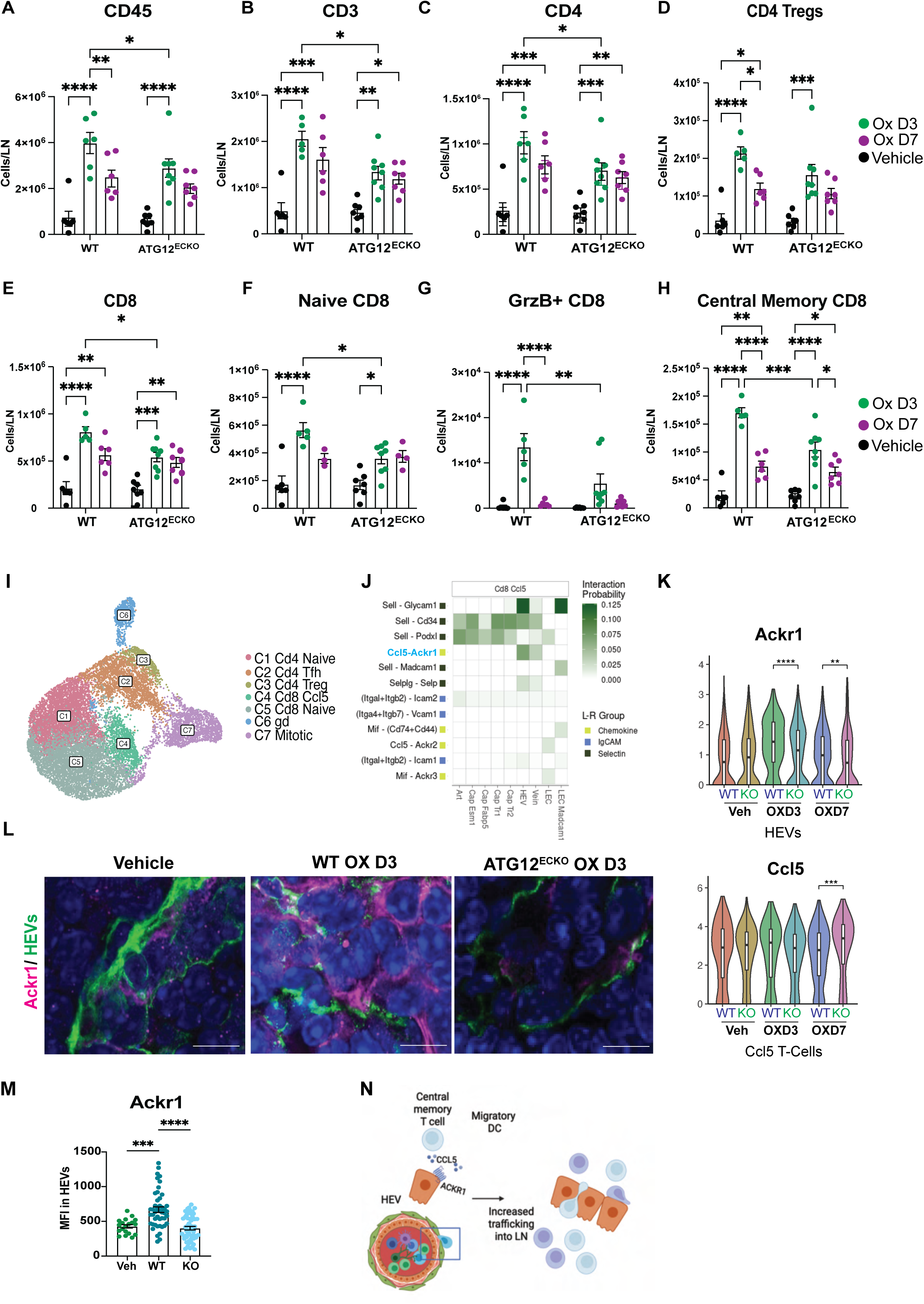
Loss of autophagy reduces HEV function. (A-H) Immune cell number characterization of mouse LNs in WT and ATG12^ECKO^ samples treated with vehicle, OX-D3 and OX-D7 by FACs. Each dot represents one mouse. (I) UMAP colored by unsupervised T-cell clustering in WT and ATG12^ECKO^ vehicle, OX-D3 and OX-D7 mouse LN. (J) Predicted Ligand (CD8 CCL5 T-cell) Receptor (EC) interactions using CellChat (K) Violin plots for Ackr1 in HEVs across samples and Ccl5 in Ccl5+ CD8 T-cells. (L) Immunofluorescence staining of HEVs (Meca-79) and Ackr1 in OX Day 3 treated LNs (M) Quantification of (N) MFI Ackr1 in HEVs. (N) Schematic representation of Ackr1-Ccl5 receptor ligand interaction effect on lymphocyte infiltration. *p < 0.05; **p < 0.01; ***p < 0.001; ****p < 0.0001

Next, by leveraging scRNAseq data from T-cells of WT and ATG12^ECKO^ homeostatic and inflamed LN samples, we employed Cellchat analysis to identify interactions between ECs and T-cells **(Figure 4I, S4C, D)**. This analysis highlighted well-studied interactions, such as SelL/Glycam1 on all T-cells, which is perturbed by the marked decrease in Glycam1 (**Figure 3C**), and likely contributes to the reduction in lymphocyte entry on OX-D3 (**Figure S4D**). Additionally, Cellchat highlighted an underexplored interaction between Ackr1 on HEVs and Ccl5 in memory T-cells (Ccl5+ Cd8+ T-cells) (**Figure 4J**). At the RNA and protein level, Ackr1 was downregulated in HEVs, while *Ccl5* was mostly unaltered on T-cells at the peak of inflammation (**Figure 4K-M**). Ackr1 has been shown to favor transcellular diapedesis^29,30^ in other settings. Reduction of Ackr1, may therefore also contribute to the reduced lymphocyte infiltration observed in ATG12^ECKO^ mice **(Figure 4N)**.

In sum, these results provide evidence that autophagy has a specific function in HEVs by ensuring PNAd+ surface molecules and fate identity. Such a mechanism is critical when HEVs need to produce more PNAd during inflammation to cope with a higher demand of lymphocyte influx and priming.

These results raise the question of how autophagy maintains the fate and function of HEVs. We first focused on the LTβR axis (**Figure 5A**), given its already known implication in HEV maintenance and our observation that the relative activity of the downstream TFs *Relb* and *Nf-kb2* were reduced in LN HEVs of ATG12^ECKO^ mice at OX-D3 and OX-D7 (**Figure 3B)**. Validating this observation, *Relb* and *Nf-kb2* activity, as well as expression of the downstream target *Ccl21a* were significantly dampened in HEVs on OX-D3 and OX-D7(**Figure 5B, C**).

**Figure 5:**
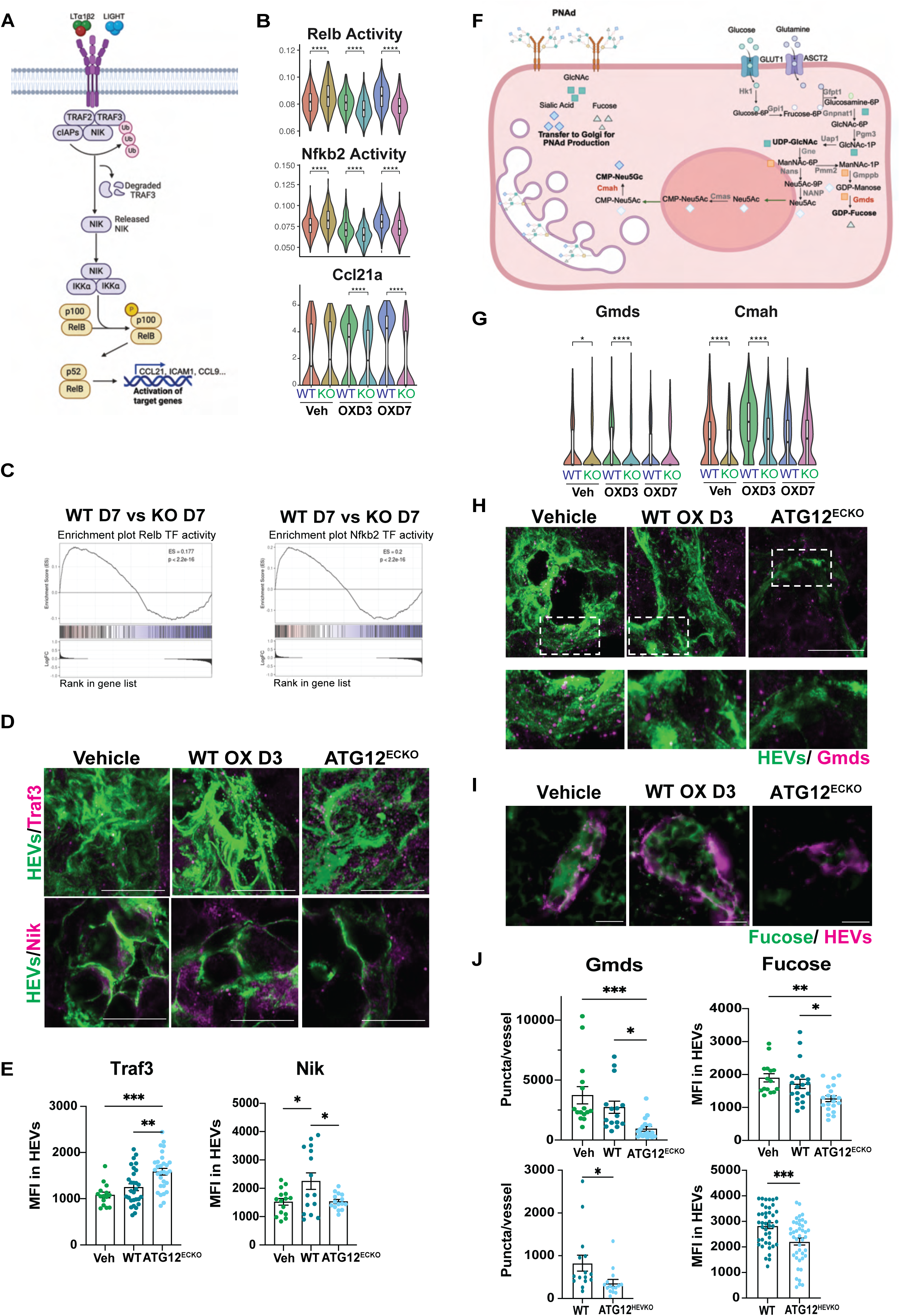
Autophagy blockade affects HEV maintenance via PNAd production. (A) Schematic representation of lymphotoxin beta receptor signaling (B) Violin plots for Relb and Nfkb2 activity, and Ccl21a expression in HEV clusters across samples. (C) GSEA plots for Relb activity (Left) and Nfkb2 activity (right) as predicted by SCENIC in WT vs ATG12^ECKO^ mice on OX D7. (D) Immunofluorescence staining of Traf3 and Nik in HEVs from Vehicle or WT vs ATG12^ECKO^mice OX D3. Scale bar=10um (E) Quantification of MFI of Traf3 and Nik in (D) (F) Schematic representation of Fucose, GlcNAc, and sialic acid production in HEVs (G) Violin plots for Cmah and Gmds expression in HEVs across samples (H) Immunofluorescence staining of Gmds in HEVs from Vehicle or WT vs ATG12^ECKO^ mice OX D3. Scale bar=10um. (I) Immunofluorescence staining of Fucose (AAL-lectin) in HEVs in from Vehicle or WT vs ATG12^ECKO^ mice OX D3. Scale bar=10um. (J) Quantification of Gmds puncta per HEV and Mean MFI of Fucose (AAL-lectin) in Vehicle or WT vs ATG12^ECKO^ mice OX D3 (Top) or WT vs ATG12^HEVKO^ mice OX D3 (Bottom). *p < 0.05; **p < 0.01; ***p < 0.001; ****p < 0.0001

Lymphocytes and DCs are the major providers of the LTβR ligands LTα2β1 and LIGHT in the LN^31–33^. Although lymphocyte levels are reduced in the LNs of ATG12^ECKO^ mice, it is unlikely that such a reduction leads to the drastic loss of LTβR signaling in ATG12-KO HEVs. Interestingly, recent evidence suggests that autophagy is involved in regulating Traf3, which inhibits LTβR signaling by targeting NIK for degradation^32,33,34^. Previous studies have shown that Traf3 can be degraded not only by ubiquitination but also via autophagy in autophagosomes^35,36^. In line with this observation, we found increased Traf3 protein levels in ATG12-KO HEVs, accompanied by a corresponding decrease in Nik levels, supporting the notion that autophagy promotes LTβR activity in HEVs (**Figure 5D, E**). Notwithstanding, while LTβR signaling is dampened, it is not entirely blocked, suggestive of additional pathways that together cause the striking phenotype observed in autophagy-deficient HEVs.

We next investigated the implication of autophagy in the proper production of PNAds, the key components and hallmarks of HEVs. As autophagy is, at its core, a recycle process to maintain cellular homeostasis^12^,we surmised whether autophagy is involved in the production of the metabolites and enzymes involved in forming PNAd chains and support HEVs to deal with inflammatory stress. Under these conditions, HEVs must expand and generate more of the fucosylated and sialylated carbohydrate structures in the ER for the modifications of the lymphocyte adhesion molecules (**Figure 3C**) and increase protein synthesis of the necessary enzymes. Indeed, we found that autophagy plays a crucial role as the expression of most enzymes, involved in PNAd biosynthesis and assembly, was downregulated in ATG12-KO HEVs **(Figure 3C**).

Among those were the enzymes *Gmds* and *Cmah* that generate cellular pools of Sialic acid (Neu5GC) or Fucose, respectively, two key components of the PNAd sugar chain, (**Figure 5F, G**). Due to reduced Gmds levels, less fucose was also observed in OX-D3 and OX-D7-treated ATG12-KO HEVs as measured by the fucose-binding AAL-lectin (**Figure 5H-J**)^37^. The decrease in the PNAd production machinery was similarly observed when in Atg12KO HEVs using the Chst4 CreER reporter, despite incomplete KO (**Figure 5J**). Thus, it is conceivable that the shortage of the necessary building blocks to produce PNAd in autophagy-deficient HEVs diminishes PNAd on the cell surface and contributes to the loss of HEV fate.

### Autophagy is linked to the UPR to cope with inflammatory HEV stress

In line with our proposition that autophagy protects HEVs from inflammatory ER stress, we noted an increased relative activity of the *Xbp1* in LN-HEVs of OX-D3 and OX-D7 mice (Figure **3B**). Xbp1 is the primary downstream target of the Ire1 branch of the UPR induced by ER proteostasis loss. The accumulation of unfolded proteins in the ER lumen drives oligomerization-mediated autophosphorylation of Ire1, activating its cytosolic RNase domain and splicing the mRNA of Xbp1 (Xbp1s). Upon translation and nuclear translocation, Xbp1s activates the transcription of several genes implicated in restoring ER homeostasis, lipid biosynthesis, and glycosylation pathways ^38^.

scRNA-seq analysis revealed that HEVs, compared to other ECs in LNs, expressed the highest levels of ER molecular chaperones, ER-associated protein degradation components, and UPR-signaling (**Figure 6A**). Strikingly, autophagy deficiency in HEVs suppressed the ER stress response in OX-D3- and OX-D7-treated ATG12^ECKO^ mice, revealing that loss of autophagy either directly or indirectly affects UPR status in HEVs (**Figure 6B-D, S5A**). Congruent with reduced Xbp1 expression and activity, we also noted that the other two branches of the UPR, Atf4 and Atf6 were also downregulated in HEV of OX-treated ATG12^ECKO^, with Atf6 downregulation noticeable at OX-D7 (**Figure 6D-F, S5B).** Reduced Xbp1s activity corresponded with a significant reduction of p-Ire1a in ATG12-KO HEVs (**Figure 6G-H**). Given that Xbp1, Relb, and Nf-kb2 are three of the top transcription factors in HEVs (**Figure 3B**), we wondered to which extent our autophagy-deficient phenotype was due to loss of LTβR signaling or loss of UPR activation. We thus used SCENIC inferences to check the regulon activities of the three TFs in relation to enzymes associated with the PNAd production on OX-D3 (**Figure 6I**). Unsurprisingly the entire regulon network was down in KO HEVs as compared to WT. While the expression of most enzymes was associated with both LTβR signaling and XBP1, sialic acid production via Cmah was only associated with LTBR signaling. In contrast, fucose production via Gmds was associated only with XBP1 activity. These data suggest that autophagy acts upstream of both LTβR signaling and XBP1 activation (**Figure 6I**). Taken together, autophagy and ER stress responses act in concert to maintain HEV fate, and both pathways likely maintain the activation of each other in HEVs (**Figure 6J**). Further support stems from a recent report, which provides evidence that genetic and pharmacological perturbations of Xbp1 in HEVs compromise PNAd production and flatten their appearance, resulting in decreased lymphocyte homing, thereby displaying a phenotype reminiscent of that in autophagy-compromised HEVs^39^.

**Figure 6:**
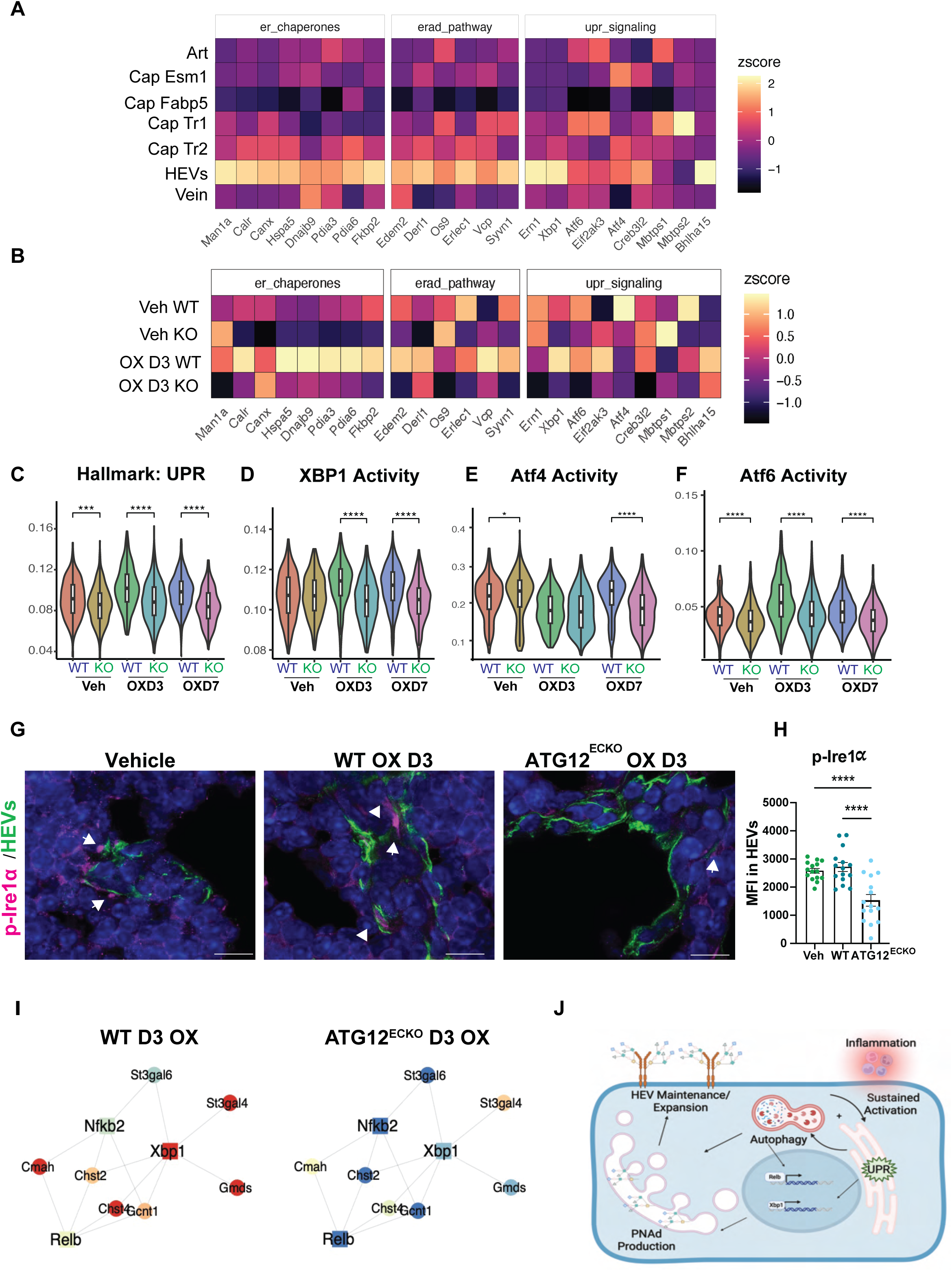
Autophagy interacts with UPR in regulating cellular stress in HEVs. (A) Heatmap of ER chaperone, ERAD, and UPR-related protein expression across EC clusters (B) Heatmap of ER chaperone, ERAD, and UPR-related protein expression in HEV cluster across Vehicle and OX D3 samples. (C) Violin plot of Hallmark UPR in HEV cluster across samples (D) Violin plot for Xbp1 activity in HEV cluster across samples (E) Violin plot for Atf4 activity in HEV cluster across samples (F) Violin plot for Atf6 activity in HEV cluster across samples (G) Immunofluorescence staining of p-Ire1α in HEVs on OX D3 in Vehicle or OX treated WT and ATG12^ECKO^ mice. Scale Bar = 10um Arrows denote positive staining. (H) Quantification of MFI of p-Ire1α in (G) (I) Gene-regulatory network (GRN) predicted by SCENIC, colored by gene expression (round nodes) or regulon activity (square nodes) in WT vs KO HEVs OX D3. (J) Proposed interaction of the UPR with autophagy in maintaining HEVs *p < 0.05; **p < 0.01; ***p < 0.001; ****p < 0.0001

### HEVs promote the inflammatory response in skin psoriasis

Consequently, autophagy-deficient HEVs become dysfunctional and lose their identity, resulting in reduced lymphocyte influx and subsequent priming. This observation raised the question of whether impairing HEV function would dampen chronically inflammatory diseases, such as autoimmune diseases. Psoriasis is a chronic, immune-mediated skin condition that is associated with an increased risk of developing cardiovascular disease and other autoimmune disorders. Patients develop scaly patches of chronically inflamed skin, characterized by the presence of aggregates of neutrophil and Th17 lymphocytes, as well as the release of inflammatory cytokines such as IL-17A, IL-17F, TNF, IL-22, and IL-6. In support of a recent report in which HEVs were detected in the skin of patients with various inflammatory skin diseases, including 6 psoriasis lesions, we identified several areas of Chst4+ HEVs in human psoriasis lesions from eight therapy-naïve patients with plaque psoriasis. Compared to non-HEV-BECs, HEVs were surrounded by a greater number of CD3 T-cell aggregates, highlighting their superior ability to transport lymphocytes (**Figure 7A-B**). In addition, we observed an increased number of LC3 puncta in HEVs compared to non-HEV BECs in psoriasis, which is reminiscent of heightened autophagy and is similar to the observations made in human tonsils during tonsilitis (**Figures 7C, 1L-M**). This made psoriasis an attractive target, since we can target both LN-HEVs and ectopic HEVs in the skin lesions, affecting both systemic and local inflammation.

**Figure 7:**
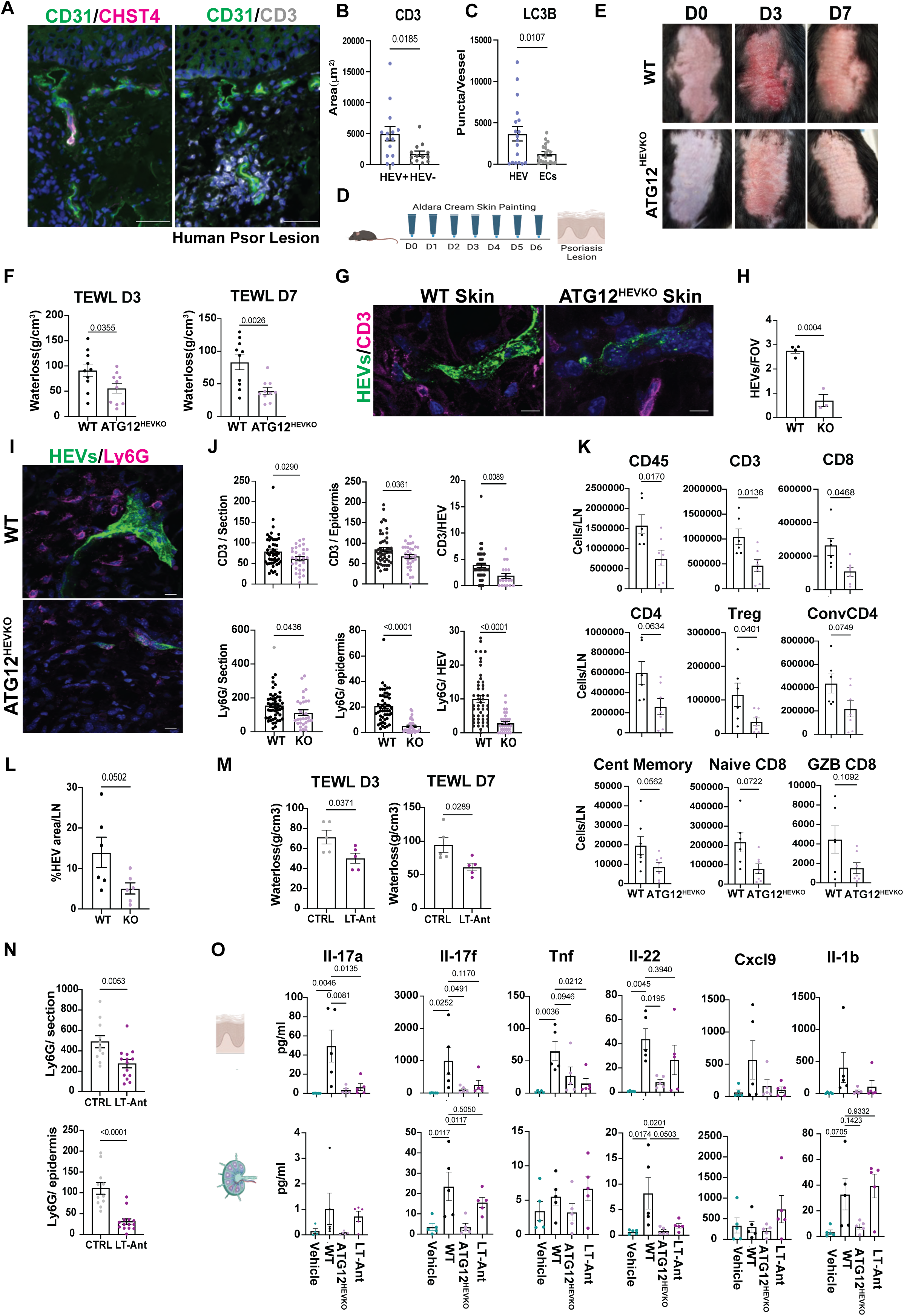
HEV blockade in psoriasis alleviates lesional phenotype. (A) Representative immunofluorescence staining (Left) of HEVs (CHST4+ CD31+) in human psoriasis biopsies, and (Right) lymphocyte/vessel staining (CD3/CD31) in the consecutive section. Scale bar=50um (B) Quantification of Lymphocyte aggregate size around HEV+ (CHST4+) and HEV-(CHST4-) vessels. (C) Quantification of LC3B puncta in HEVs (CHST4+ vessels) vs BECs (CHST4-vessels) in human psoriasis biopsies from n=8 patients. (D) Schematic of aldara (imiquimod) treatment schedule (E) Images of psoriasis lesion in WT and ATG12^HEVKO^ mice on D0, D3, and D7. (F) Trans-epidermal water loss (grams/cm^3^) on D3 and D7 aldara treatment in WT and ATG12^HEVKO^ mice. (G) Representative images of HEVs and CD3 (lymphocytes) in skin lesions from WT and ATG12 ^HEVKO^ mice. Scale bar= 10um (H) Quantification of the average number of HEVs per FOV in skin lesions from WT and ATG12 ^HEVKO^ mice. Each dot represents one mouse. (I) Representative images of HEVs and Ly6G (neutrophils) in skin lesions from WT and ATG12 ^HEVKO^ mice. Scale bar =10um (J) Quantification of lymphocyte (top) and Neutrophil (bottom) numbers per section (left), per epidermis section (middle), or per area (within 50 um) around HEVs (right). (K) Immune cell number characterization of mouse LNs from WT and ATG12 ^HEVKO^ by FACs. Each dot represents one mouse. (L) Quantification of HEV area per LN in WT vs ATG12 ^HEVKO^ (M) Trans-epidermal water loss (Grams/cm^3^) on Day 3 and Day 7 aldara treatment in control and LTΒR-ANT treated mice (N) Quantification of Neutrophil numbers per section (left) and per epidermis section (right) (O) Quantification of cytokine amount (pg/ml) in skin samples (top) and LN samples (bottom) for Vehicle, WT, ATG12 ^HEVKO^, and LTΒR-ANT treated mice

To test this hypothesis, we utilized the well-established Aldara (imiquimod) mouse model of psoriasis, reminiscent of human disease, in which imiquimod cream is administered to the skin for 7 consecutive days This induces reddening and plaque formation of the skin, concomitant with increased transepidermal water loss (TEWL)(**Figure 7D**)^40^. We then induced psoriasis WT-mice and ATG12^HEVko^ mice, in which ATG12 is deleted in Chst4+ HEVs, to elicit the differences in skin and LN inflammation. Notably, ATG12^HEVKO^ mice displayed a less severe skin rash after 3 and 7 days and less transepidermal water loss in the skin (**Figure 7E, F**), suggestive of a reduced inflammatory response when autophagy was impaired in HEVs, by comparison TEWL measurements in healthy skin ranged from 2-12 g/cm^3^. Immunohistochemical analysis of the skin lesions revealed HEVs in psoriatic skin (Ps-HEVs) lesions of WT-mice, similar to the observations in human psoriasis. However, HEVs were hardly detectable in lesions of ATG12^HEVKO^ mice, and mostly as single cells (**Figure 7G-I**). These results suggest that autophagy may also affect the formation of ectopic HEVs during inflammation in the skin. Consequently, while we found a substantial accumulation of Ly6G+ neutrophils and CD3+ T-cells in both the epidermal and dermal layers of psoriatic lesions, these immune infiltrates were significantly reduced when autophagy was impaired in HEVs (**Figure 7G,I,J**). Ps-HEVs of WT mice were surrounded by multiple immune cell aggregates, while the few HEV remnants in the lesions of ATG12^HEVKO^ mice yielded small clusters of neutrophils and lymphocytes. Of note, neutrophils in psoriasis of ATG12^HEVKO^ mice appeared to remain deep within the dermis, while neutrophils in WT lesions were abundantly found in both the dermis and epidermis (**Figure S6A**).

To understand the extent to which the diminished immune infiltrates in psoriasis are a consequence of ATG12KO HEVs in skin versus LNs, we interrogated the immune infiltrates of LNs harboring autophagy-proficient or deficient HEVs in psoriasis-bearing mice. As anticipated from the results obtained from LNs of ATG12^ECKO^ mice during oxazolone-induced inflammation (**Figure 4A-H**), we also observed reduced infiltration of the different CD4 and CD8 T-cell populations in LNs of ATG12^HEVKO^ psoriasis-bearing mice (**Figure 7K**). This was accompanied by and a result of the reduced HEV area, lower PNAd levels, and lack of upregulation of inflammation-induced leukocyte adhesion molecules such as P-selectin in these LNs (**Figure 7L and S6B, C**). These results imply that impairing HEVs in LN and skin leads to overall reduced infiltration and activity in psoriatic inflammation. It is conceivable that by reducing the influx of lymphocytes in the LN, fewer lymphocytes will leave the LN and enter the circulation. In support, we observed smaller spleens with reduced weight and length, reflecting fewer lymphocyte infiltrates **(Figure S6D).**

Given the robust attenuating effects on the inflamed psoriasis phenotype by solely dampening HEV function, we sought to test the therapeutic potential of inhibiting HEVs in psoriasis by blocking LTβR signaling. The LTβR pathway is implicated in autophagy-dependent maintenance of HEVs (**Figure 5**), and is known to be essential for HEV formation and maintenance^4,15,27,41^. Using an LTβR antagonist (LTβR-ANT) in psoriasis-bearing mice, we obtained similar results to those using a genetic knockout of ATG12 in HEVs in terms of reduced rash **(Figure S6E)**, transepidermal water loss **(Figure 7M)** and neutrophil and lymphocyte infiltration in the skin **(Figure 7N, S6F).** Furthermore, HEV area and lymphocyte presence in the LN were similarly reduced (**Figure S6G, H).**

The formation of psoriasis lesions relies on complex cytokine/chemokine signaling. DCs produce TNF and IL-6, which promote the differentiation of naïve T-cells into Th17 cells^42^. Th17 cells are considered the major actors that secrete Il-17. Both Il-17a and Il-17f are well accepted to play a role in the pathogenesis of psoriasis^43^, as well as Il-22, which is also released by T-cells and contributes to pathogenesis^44^. Il-17 then activates keratinocytes, which produce several chemokines/cytokines, including Cxcl9, Cxcl1, and Il-1β, to recruit leucocytes (including neutrophils and Th17 cells) to the site of inflammation^42,43^.

To understand of how impaired HEV function by either abrogating autophagy or LTβR signaling in the psoriasis mouse models affected the release and effects of these inflammatory cytokines, we performed a 48-cytokine/chemokine panel analysis using the OLink platform in vehicle, WT, ATG12^HEVKO^, and anti-LTΒR treated psoriasis mice (**Figure 7O, S6I**). The most prominent cytokines Il-17a, Il-17f and Il-22 were all significantly reduced in the skin ATG12^HEVKO^ mice, followed by a similar trend of reduced Tnf, Cxcl9, and Il-1β levels (**Figure 7O**). Similar results were obtained from the cytokines of LTβR-ANT-treated skin. Il-17a and Tnf were significantly reduced with a trend of reduced Il-17f, Cxcl9 and il-1β (**Figure 7O**). Interestingly, we observed the same reduction of all these cytokines in the LNs of ATG12^HEVKO^ or LTβR-ANT-treated psoriasis-bearing mice, albeit the overall levels were substantially lower than in the skin. Moreover, Il-6, Cxcl1 and G-CSF were significantly reduced in the skin of Autophagy HEV-KO and LTΒR-ANT treated mice, which may contribute to the marked decrease we see in neutrophil infiltration (**Figure S6I**). Together, these data show the therapeutic potential of targeting HEVs in autoimmunity.

## Discussion

HEVs are essential for proper immunological responses as they act as conduits for lymphocyte infiltration in SLOs and at sites of local inflammation. Here, we demonstrate that autophagy plays a pivotal role in HEVs, distinct from that of other ECs. Its activity in HEVs is heightened during inflammation, to ensure the maintenance of HEV fate and function. HEVs are defined by their coat of surface PNAds, decorated with glycan structures, including the 6-sulfo sialyl Lewis X moieties, which serve as L-selectin ligands for the efficient influx of naïve lymphocytes. During inflammation, HEVs must expand and increase PNAd biosynthesis to accommodate the heightened demand for lymphocyte infiltration and priming, thereby combating infections. It is thus not surprising that HEVs contain a prominent ER and Golgi network to enable the complex biosynthesis of these structures. We provide several lines of evidence that the autophagy program ensures the production of the PNAd carbohydrate structures to maintain HEV fate via the LTβR signaling and the UPR. Genetic perturbation of autophagy in LN HEVs and ectopic HEVs in psoriasis lesions diminishes PNAd levels and evokes a flattened appearance of HEVs concomitant with signs of transdifferentiation into capillary endothelial cells. Consequently, lymphocyte influx is severely restrained. The effects are most prominently seen during inflammation, but a lack of autophagy also shows signs of dedifferentiation in homeostatic HEVs, albeit within several weeks, due to their low turnover rate.

Autophagy maintains HEVs in part by degrading Traf3^35^, a negative regulator of LTβR, the essential HEV signaling pathway ^4,8,27,45^, as well as by providing metabolic building blocks necessary to produce PNAd (Fucose, Sialic acid). Congruently, it has already been shown that sialyloligosaccarides can be broken down by autophagy^46^. In addition to these metabolites, O-glycans are essential for producing a functional PNAd molecule. While not investigated in our study, O-GlycNAylation, like autophagy, is activated by nutrient availability cues and interacts with the autophagy pathway at different levels. O-GlcNAc can specifically activate the essential autophagy protein ULK1, whereas SNAP29, a SNARE required for autophagosome/lysosome fusion, becomes O-GlcNAcylated in the absence of autophagy^47,48^.

A specific feature of HEVs is that they exhibit heightened UPR and ER-related pathway activity in comparison to other BECs. This is likely due to the increased need for protein folding/membrane trafficking of PNAd-related mucins and enzymes. In HEVs, lack of autophagy leads to a downregulation of the UPR, and more specifically of the transcription factor Xbp1, which was recently shown to be directly involved in the production of PNAd through the transcriptional expression of the conjugating enzymes that add Fucose, sialic acid, and GlyNAC to HEV sialomucins ^39^.

Autophagy-deficient HEVs, like Xbp1-knockout HEVs^39^, exhibit a substantial reduction in the essential enzymes for PNAd production. Therefore, in addition to directly affecting LTβR signaling and PNAd metabolite production, defective HEV function upon autophagy blockade may be exacerbated by the interaction with the UPR system. Several studies have delineated mechanisms through which UPR and autophagy functionally interact; ATF4 specifically binds to the promoter of many ATG-related genes^49^. IRE-1 activation via Traf2 has also been shown to stimulate autophagy directly via a Beclin1-dependent mechanism^50^. While most of the current literature suggests the UPR is upstream of autophagy activation in response to stress^28^, the interactions between the two pathways may not be hierarchical in HEVs, but it appears that they rather operate in a continuous feedback loop, where both pathways keep the other stimulated as an adaptation mechanism against inflammatory stress, and produce more PNAd, to maintain HEV status. Another line of support that autophagy and UPR are required to maintain HEV fate stems from pancreatic beta cells^51^. These cells have a constitutive basal level of UPR, like HEVs, which is increased during obesity-related stress. Loss of autophagy in pancreatic beta cells has been shown to compromise the UPR machinery by downregulating the expression of diverse UPR-associated genes. Consequently, proinsulin oxidative folding in the ER was disrupted and proinsulin trafficking from the ER to Golgi impaired, leading to decreased insulin production and a loss of the pancreatic beta cell fate^5253^. Likewise, B-cells upregulate Xbp1 during plasma cell differentiation^54^. Therefore, in addition to maintaining cellular homeostasis, autophagy and UPR can regulate cell identity, by acting non-hierarchically, depending on the cellular context and stress signal. As defective portions of the ER are specifically degraded by autophagy, through a process known as ER-phagy, to restore organelle function^55^, it is intriguing to speculate that the lack of ER maintenance by autophagy could contribute to this phenotype.

Evidently, the Xbp1 knockout in HEVs^39^ resembled the phenotype of autophagy-deficient HEVs; i.e., compromised PNAd production, flattened morphology, and reduced ability for lymphocyte homing. Consequently, autophagy deficiency led to a loss of the HEV cell fate and differentiation into capillary endothelial cells in the LNs. Interestingly, we observed that ectopic intratumoral HEVs arise from postcapillary venules and also differentiate back upon loss of LTβR signaling^8^. These data indicate that autophagy is an essential mechanism for maintaining the cell fate of HEVs.

It is important to note that autophagy affects immune cell trafficking in an endothelial subtype-specific manner. Our previous work, studying autophagy in LN lymphatic ECs, found an autophagy-specific mechanism for the production of sphingosine-1-phosphate, the key lipid mediator that guides lymphocyte egress from the LN into the circulation ^41^. Here, we revealed that autophagy-compromised HEVs exhibit a defect in regulating lymphocyte influx, it highlights the importance of EC autophagy for the proper trafficking of immune cells. In contrast, the lack of autophagy in BECs during inflammation enhanced leukocyte adhesion molecule expression and increased neutrophil trafficking across EC^14^. Our study of tumor BECs provided similar results, where lack of autophagy in tumor BECs led to increased adhesion molecule expression and consequent increased T-cell infiltration in the tumor^15^. Therefore, while EC autophagy is clearly necessary for proper EC-immune interactions, its role is very distinctive depending on which vascular bed is targeted.

Treating autoimmune diseases remains challenging due to the adverse side effects of directly targeting the immune system. Psoriasis is an autoimmune disease that affects approximately 8 million Americans and 125 million people worldwide^56^. Recent observations and our studies have shown that ectopically induced HEVs can develop within psoriatic lesions^10^. Given that HEVs spur leukocyte influx, could HEV blockade prove to be an effective psoriasis therapy? Although various treatments are available for moderate to severe psoriasis, the most effective therapies (IL-17 and IL-23 blockers, and TNF-alpha inhibitors) can cause severe side effects in patients, including an increased risk of moderate to severe infections due to their systemic targeting of the host immune system^57^. Therefore, an indirect vascular-targeting approach could offer a promising way to preserve immune function while reducing the risk of autoimmunity. We found that inhibiting HEV function by blocking autophagy, significantly attenuated the inflammatory severity of the disease, as transepidermal water loss, cytokine production, and the presence of lymphocytes and neutrophils in the lesional skin were significantly reduced. As pharmacological perturbations of EC-targeted autophagy would not be translationally effective, due to the promiscuous effects of autophagy in different vascular compartments^14,15,41^, we directly blocked LN and ectopic HEVs in psoriasis lesions with an LTβR-antagonist ^27,45^. The inflammation-blocking effects of the LTβR antagonist in psoriasis-bearing mice were comparable to those when autophagy was genetically deleted in HEVs. These results are encouraging because blocking HEV function reduces both local and systemic inflammation. Fewer naïve T cells enter the lymph node, resulting in fewer primed T cells circulating in the blood. Conversely, altered HEV formation in the skin leads to reduced T-cell entry, subsequently decreasing T_H17_ differentiation and diminishing downstream neutrophil activation. It is intriguing to speculate that the approach of blocking HEV function in LN and at local sites of HEV/TLS-containing chronic inflammatory and autoimmune diseases^58^ may be specifically attractive, as it tones down inflammation without compromising immune function. Therefore, HEV-blocking therapies may present a new avenue for exploring the treatment of various autoimmune diseases.

## RESOURCE AVAILABILITY

### Lead Contact

further information and requests for resources and reagents should be directed to and will be fulfilled by the lead contact, Gabriele Bergers

### Materials availability

Reagents generated in this study will be made available on request, but we may require a payment and/or a completed Materials Transfer Agreement if there is potential for commercial application.

### Data and Code availability

For scRNA-seq, all murine raw and processed sequencing data are available in GEO under accession number GSE300153 and will be publicly available as of the date of publication. Accession and identification numbers are listed in the Key resources table

### Experimental Models and subject details

#### Animal strains

Animal procedures were approved by the Institutional Animal Care and Research Advisory Committee of the KU Leuven (ECD 078/ 2021) and were performed following the institutional and national guidelines and regulations. C57BL/6 mice were purchased from Charles River or obtained from the KU Leuven animal facility.

All conditional knock out mice in this study were of a C57BL/6 background.

Conditional ATG12 EC knockout mice (Cdh5-Cre^ER^x ATG12_lox/lox_ RFP mice) were generated by intercrossing ATG12_lox/lox_RFP^59^ mice with the tamoxifen-inducible, endothelial cell-specific Cdh5-CreER^60^ mouse line. Cdh5-CreERnegxATG12_lox/lox_ RFP mice littermates were used as controls.

Conditional ATG7 EC knockout mice (Cdh5-Cre^ER^x ATG7_lox/lox_ mice) were generated by intercrossing ATG7_lox/lox_ mice with the tamoxifen-inducible, endothelial cell-specific Cdh5-CreER mouse line. Cdh5-CreERnegxATG7_lox/lox_ mice littermates were used as controls.

Chst4-tdT reporter mice (C57BL/6J) were generated by intercrossing tamoxifen-inducible Chst4-Cre(ER)T2/iRFP mice^8^ with Rosa26-LSL-tdTomato reporter mice.

Conditional HEV specific ATG12 knockout mice (Chst4-Cre^ER^x ATG12_lox/lox_) were generated by crossing Chst4-Cre (ER)T2/iRFP mice with ATG12_lox/lox_ RFP mice. For tracer experiments, CHST4-tdT reporter mice were used as controls. For all other experiments, CHST4-CRE^ER^negxATG12_lox/lox_ RFP were used as controls.

RFP-GFP-LC3 mice were purchased from Jackson (#027139)^21,22^.

For all experiments, animals were maintained in individually ventilated cages in a room with controlled temperature and humidity under a 12-h light/12-h dark cycle with ad libitum access to water and food. Information on age and sex of mice can be found in the ’method details’-section.

#### Human samples

The collection of human tonsilitis samples was done under the license of the Commission d’Ethique Biomédicale Hospitalo-Facultaire, Brussels, Belgium, references 2021/03JUI/264, 2019/17JUI/261 and 2020/29JUI/345. The collection of human psoriasis samples was done under the license 19–8636-BO (Essen, Germany). Written informed consent for donating tissue was obtained from each patient. The samples were kept anonymously and handled according to the ethical guidelines set by the University of Turku.

#### Mouse Trials

##### Steady-state

7–10-week-old male and female C57BL/6 ATG12 fl/fl xCDH5-CreER mice were treated with tamoxifen 5 times according to the Jackson protocol. Mice were left for 6 weeks before analysis.

##### Oxazolone

7–10-week-old male and female C57BL/6 mice (LC3B-RFP-GFP, ATG12 fl/fl xCDH5-CreER, ATG7 fl/fll xCDH5-CreER, CHST4-CreER Tomato or ATG12 fl/fl xCHST4-CreER) mice were treated with tamoxifen 5 times according to the Jackson protocol. Mice were shaved with Veet on the abdomen and 50 ul of 4% Oxazolone (3:1 acetone: olive oil) was applied to each area. Mice were left for 3 to 7 days depending on the experiment.

##### Psoriasis

7–10-week-old male and female ATG12 fl/fl xCHST4-CreER mice were pretreated once with tamoxifen, and their backs were shaved with an electric shaver, followed by Veet. A fourth of a packet of Aldara cream (imiquimod 5%) was applied to the back of mice daily in an even layer from day 0 to day 6. Tamoxifen was administered every other day during treatment.

*LTBR-antagonist*: C57BL/6 mice were shaved on their backs with an electric shaver, followed by Veet. A fourth of a packet of Aldara cream (imiquimod 5%) was applied in an even layer daily from day 0 to day 6. 2mg/kg of LTβR-antagonist (Tebu bio, #71122), was administered intraperitoneally on day 1 and 4 post-Aldara cream administration.

Trans epidermal water loss was measured with a Tewameter TM Nano monitor (Courage and Khazaka).

##### Intravital imaging

Prior to imaging, mice were intravenously injected with MECA-79-AF488 (20ug). WT mice were also injected with a CD31-AF594 (20ug). Mice were anaesthetized under 2% isoflurane/air mixture. An imaging window was placed on top on the inguinal lymph node previously established for mammary gland imaging^59^. Mice were placed on an imaging box and kept at 32°C during imaging. Inguinal lymph node imaging was performed on an inverted Leica SP8 Dive system (Leica Microsystems) equipped with 3 tunable hybrid detectors, a MaiTai eHP DeepSee laser (Spectra-Physics) and an InSight X3 laser (Spectra-Physics) using the Leica Application Suite X (LAS X) software. All images were collected at 12 bit, acquired with a 25 x water immersion objective with a free working distance of 2.40 mm (HC FLUOTAR L 25x/0.95 W VISIR 0.17) and a z-step of 10um. AF488 and AF594 were excited with a 780nm laser. Analysis of the timepoints was performed with ImageJ.

### Tissue dissociation and sample preparation

For all EC analysis (FACs+ scRNAseq) a previously published protocol was utilized^62^. Briefly, lymph nodes were extracted and minced, then incubated at 37°C in serum-free RPMI containing (0.8mg/ml Dispase II, 0.2 mg/ml Collagenase P, and 0.1mg/ml Dnase I) for 15 minutes, refreshing twice with manual dissociation between each incubation. Remaining enzymes were quenched in MACS buffer (Miltenyi), and suspensions were passed through 70 μm strainers to remove any fat pieces. For EC flow cytometry analysis, immune cells were depleted using the positive selection mouse CD45 microbeads (Miltenyi). The negative fraction that contained the remaining endothelial cells was collected and washed with PBS before flow cytometry staining.

For immunoprofiling of LNs, LNs were squashed on a 70um cell strainer and rinsed with RPMI media prior to cell staining.

### Flow Cytometry

Single cell suspensions were blocked with anti-CD16/32 blocking antibodies (BD Biosciences), and then stained for 30 minutes on ice with antibody cocktail containing: CD31 (clone 390), MECA-79 (clone MECA-79), CD3 (clone 17A2), CD4 (clone GK1.5; clone RM4-5), CD8α (clone 53–6.7), CD11c (clone N418), CD19 (clone 1D3;B4), CD45 (clone 30-F11), CD62L (clone MEL-14), CD44 (IM7), Foxp3 (clone REA788), Granzyme B (clone NGZB; clone QA16A02), Ly6C (clone HK1.4), MHC-II (clone M5/114.15.2), Ki67 (clone 16A8). Dead cells were excluded using the Fixable Viability Dyes (eBioscience, Biolegend) or 7AAD (eBioscience). Cells were fixed with Fixation buffer and permeabilized by Foxp3/Transcription Factor Staining Buffer from (ThermoFisher) before intracellular staining. Counting beads were added prior to sample run at the cytometer.

For EDU experiment, mice were pretreated interperitoneally with 250 ug/mouse for 3 consecutive days with EDU. Following surface marker staining. Click-it EDU reaction was performed according to the manufacturers protocol (Thermo scientific)

Data for immunoprofiling or EC profiling were acquired on a BD FACSFortessa (BD Biosciences). Samples for scRNA-seq were sorted by a BD FACs Aria Fusion (BD Biosciences). Results were analyzed with FlowJo version 10.8.0 (Becton Dickinson, Ashland).

### Immunostaining

Excised LNs and skin were fixed in 2 % paraformaldehyde (PFA) at 4C overnight, followed by 30% sucrose at 4C overnight before being embedded in OCT (Leica). For the preparation of agarose sections, tissues were fixed in 2% PFA at 4C for 24h, then embedded in 4% low melting agarose (in PBS).

10μM thick cryosections were stained with anti-MECA-79, anti-CD3 (17a2), anti-Ly6G (1A8), anti-CD31 (polyclonal, or 2H8), anti-P-selectin (RB40.34), anti-Gmds (polyclonal), anti-Ackr1 (polyclonal), anti-Nik (polyclonal), anti-Traf3 (polyclonal), anti-Madcam1 (Meca-367), anti-4-1BB (E2J5H), anti-Vwf (polyclonal), anti-Ccl21a (Polyclonal), anti-Icam1 (3E2), anti-Vcam1 (429), anti-Lc3B(polyclonal), AAL-Lectin (vector labs), anti-p-Ire1 (polyclonal). Alexa-fluorophore conjugated secondary antibodies were applied when primaries were unconjugated, nuclei were counterstained with Dapi and mounted in Dako mounting medium. Images were taken using an Observer z1 microscope (Zeiss) linked to an AxioCam MRM camera (Zeiss) with 20X, 40X, and 63X objectives or Zeiss LSM 880 Airyscan (Bioimaging core VIB-KU Leuven) with 63X objective.

50μM vibratome sections were stained with Meca-79. Alexa-fluorophore conjugated secondary antibodies were applied when primaries were unconjugated, nuclei were counterstained with Dapi and mounted in Dako mounting medium and imaged at a Leica Confocal SP8 with objectives of 20X or 40X.

For whole lymph node analysis used in tracer experiment, and steady-state (6 weeks) experiment as well as for Skin HEV analysis in mice were injected with 20ug Meca-79-alexa488 30 minutes prior to tissue collection. Skin was processed further as cryosections as described above. Lymph nodes were fixed for 2 hrs and imaged between two coverslips at a Leica Confocal SP8 with objectives of 20X or 40X.

Human tissue biopsies were received snap-frozen and subsequently embedded in Tissue-Tek® O.C.T compound. The embedded samples were sectioned using a cryostat, primarily at a thickness of 5 µM. The tissue sections were mounted onto SuperFrost® microscope slides (R. Langenbrinck GmbH, 03-0060) and stored at -80°C until further analysis.

Fresh frozen Human tonsil tissue and human psoriatic skin tissue were post-fixed for 10 minutes in 4% PFA. Cryosections were stained with the following primary antibodies: Tonsils: anti-MECA-79, anti-CD31(JC/70A), and anti-LC3B(D11); Skin: anti-CHST4 (polyclonal), anti-CD31 (Polyclonal), anti-CD3 (polyclonal), LC3I (5F10). Alexa-fluorophore conjugated secondary antibodies were applied when primaries were unconjugated, nuclei were counterstained with Dapi. Human skin samples were treated with Sudan Black 20 minutes post staining to remove the background in the tissue. All samples were mounted in Dako mounting medium.

Images were taken using an Observer z1 microscope (Zeiss) linked to an AxioCam MRM camera (Zeiss) with 20X, 40X, and 63X objectives or Zeiss LSM 880 Airyscan (Bioimaging core VIB-KU Leuven) with 63X objective.

Image analysis and quantitation were performed using ImageJ software version 2.0.0.

### Single-cell sequencing

#### Mouse analysis

After sorting, the single-cell suspensions were converted to barcoded scRNA-seq libraries using the Chromium Single Cell 30 Library, Gel Bead & Multiplex Kit, and Chip Kit (10x Genomics), aiming for 10,000 cells per library. Samples were processed using kits pertaining to V2 barcoding chemistry of 10x Genomics. Single samples were always processed in a single well of a PCR plate, allowing all cells from a sample to be treated with the same master mix and in the same reaction vessel. For each experiment, all samples were processed in parallel in the same thermal cycler.

To investigate the impact of autophagy deficiency on endothelial cells (ECs) during inflammation, we performed single-cell RNA sequencing on a total of 12 samples, including both CD45^-^CD31^+^ ECs and CD45^+^ immune cells, from ATG12 Cre^-^ and Cre^+^ mice under three conditions: Vehicle, OX D3, and OX D7.

Raw sequencing data was processed using CellRanger (version 7.0.1) and aligned to the mouse reference genome mm10-2020-A. Initial quality control was performed to retain cells with more than 1000 detected genes and less than 10% mitochondrial content. Doublets were identified and removed using scDblFinder. After quality filtering, a total of 62,441 cells remained for downstream analysis.

Data integration and clustering were performed using the Seurat package^63^ (version 5.0.2) in R (version 4.4). The filtered data was processed using NormalizeData, FindVariableFeatures, and ScaleData functions. Dimensionality reduction was first performed using RunPCA. Batch effects between different experimental conditions were then corrected using Harmony (version 1.2)^64^ integration based on the PCA results. Visualization was achieved using RunUMAP, and unsupervised clustering was performed using FindNeighbors and FindClusters based on the Harmony embeddings with dims = 1:15 and resolution = 0.5.

For detailed analysis of cell populations, we further subclustered major cell types separately, following the same Seurat workflow but with different dimensional and resolution parameters:

- Endothelial cells (22,272 cells): dims = 1:8, resolution = 0.4
- Myeloid cells (3,345 cells): dims = 1:6, resolution = 0.2
- Fibroblasts (8,105 cells): dims = 1:6, resolution = 0.35
- T cells (16,188 cells): dims = 1:8, resolution = 0.4
- B cells (11,280 cells): dims = 1:6, resolution = 0.2

Cell type annotation was based on the expression of canonical markers as shown in the respective figures (EC markers in **Figure S3A**, T cell markers in **Figure S4C**,).

Transcription factor (TF) activity analysis was performed using pySCENIC^65^ (version 0.9.19, using nextflow pipeline with aertslab-pyscenic-0.9.19.img container image) to identify key regulators and their target genes in different cell populations. This enabled us to characterize the transcriptional changes in HEVs following autophagy deficiency.

For trajectory analysis, we excluded lymphatic endothelial cells (LECs) and mitotic cells from the endothelial cell population. We calculated diffusion components using Palantir^66^ (version 1.3.0) in Python and visualized the results using tSNE embedding. RNA velocity analysis was performed using scVelo (version 0.3.0)^67^ with velocity vectors projected onto the tSNE plot to indicate differentiation directionality. The differentiation potential of each EC cluster was assessed using CytoTRACE (version 0.3.3)^68^.

Differential gene expression analysis between conditions was performed and visualized using the waterfall plot function from the SeuratExtend package (version 1.1)^69^, which automatically calculates differentially expressed genes between specified groups.

Pathway analysis was conducted using gene set enrichment analysis (GSEA) with hallmark gene sets from MSigDB and custom gene signatures for autophagy (macroautophagy from Reactome) and UPR activation. Enrichment scores were calculated and compared between different cell populations and experimental conditions.

Cell-cell communication analysis was performed using CellChat (version 2.1.2)^70^ to infer potential ligand-receptor interactions between HEVs and other cell types, with particular focus on interactions disrupted in autophagy-deficient conditions.

All visualizations including violin plots, heatmaps, UMAPs, waterfall plots, and GSEA plots were generated using the SeuratExtend package to ensure consistency in data presentation. Statistical significance was determined using the ggpubr package in R with default parameters, with p < 0.05 considered significant.

#### Human Tonsilitis

The scRNA-seq data from 5 healthy (patients with obstructive sleep apnea) and 4 tonsillitis patients was obtained from De Martin et al^71^, where the data was provided as a pre-filtered and integrated seurat^72^ object. The gene names were changed from Ensembl IDs to Gene symbols for easy recognition and downstream analysis. The annotations named ‘labels’ in the dataset were used to extract Endothelial cells (ECs) from the data. This annotation was verified by checking for expression of EC-related canonical markers PECAM1, CDH5, and PROX1, to distinguish between ECs and all other cell populations. The clusters with ‘labels’ BEC_1, BEC_2, BEC_3, and LEC_4 were thus extracted for further processing.

BEC_1 were identified as Pericytes, expressing classical markers PDGFRB, NOTCH3 and TPM2. LEC_4 were characterized by lymphatic EC markers PROX1 and LYVE1. These were removed and the remaining cells were processed further to identify vascular EC subtypes. BEC_2 and BEC_3 are blood endothelial cell clusters (BEC).

Batch effects were removed from the extracted ECs using Harmony integration^64^. Log-normalization using the NormalizeData function from seurat was performed on the resulting filtered EC object. This was followed by selection of highly variable genes using FindVariableFeatures and auto-scaling using the ScaleData function in seurat, with default parameters. Principal component analysis was performed for dimensionality reduction using the default RunPCA seurat function. Finally, RunUMAP function was used for data visualization, which implements uniform manifold approximation and projection (UMAP) (dims = 30) for data visualization. To find clusters, unsupervised clustering was performed in seurat using FindNeighbors followed by FindClusters function (dims = 30, resolution = 0.3).

### Assigning Cluster Annotations

The differentially expressed genes were found for each cluster using FindAllMarkers and were annotated into Arterial, Capillary, post capillary venule (pcv) and Venous cells using canonical markers (Arterial-GJA4, SEMA3G; Capillary-SPARC, EMCN, RGCC; pcv-SOCS3, C2CD4B, and venous-ACKR1, NR2F2, SELE). Capillary ECs were further subdivided into 2 subclusters-Capillary and VWF_CDH5_low_Cap (expression low levels of VWF and CDH5), based on marker genes reported in literature^26,73^.

To find HEVs, cells expressing HEV-specific markers CHST4, FUT7, GCNT1, CD24, CH25H, ENPP6, and TMEM176A were selected out of the Venous and PCV clusters. CHST4, FUT7, and GCNT1 are enzymes required for the synthesis of the 6-sulfo sLeX epitope, while the rest of the genes have previously been shown as lymph node HEV markers in tumor^8^. For this analysis, an upset plot using the genes mentioned above was drawn using UpsetR library. The cells expressing at least 6 of these genes were annotated as HEV. This strict selection ensures HEV annotation with confidence.

All violin plots, heatmaps, and feature plots were drawn using standard plotting functions from R, Seurat, and SeuratExtend package^69^.

### Comparison of Autophagy signature enrichment

To check for the expression of autophagy pathway, signatures from Bordi et al.^23^and macroautophagy from reactome^74^ database were used. The signature curated by Bordi et al. is a collection of genes that are upregulated when autophagy in cells is upregulated. First, Scran normalization^75^ was applied to the data. The expression of autophagy pathway was compared between HEV vs BECs (except Arterial cells), as well as between HEVs from non-inflamed vs tonsillitis samples using Bioconductor package AUCell^65^ enrichment. An enrichment plot for gene set enrichment analysis (GSEA) for the above signature was also performed.

### OLINK Cytokine Analysis

Tissue lysates were prepared from flash-frozen skin and lymph node samples using Bio-plex cell lysis buffer in a tissue homogenizer. Cytokine quantification was performed using Olink Mouse Target 48 Cytokine panel (Olink service platform, KU Leuven). Briefly, lysates were incubated at 4 degrees for 20h with antibodies coupled molecular probes. Molecular probes were then amplified by PCR using the T100 thermocycler, followed by quantification by qPCR in Olink Signature Q100. Quality control of the data was performed using the Olink NPX Signature software (V 1.12.1).

### Quantification and Statistical analyses

Data entry and all analyses were performed in a blinded fashion. Bar Graphs show mean value +/-SEM. Unpaired student’s t-test (two-tailed) was used for the comparison of two groups. For all bar graphs with three conditions, a one-way ANOVA with Sidak post-test was used. For Figure 3 FACs analysis, a two-way ANOVA with Tukey’s post-test was used to compare multiple groups. For single-cell analyses the statistical test used was the Wilcoxon signed-rank test, with the adjusted p-value using Holm correction. p values < 0.05 were considered significant (^∗^: p < 0.05; ^∗∗^: p < 0.01; ^∗∗∗^: p < 0.001; ^∗∗∗∗^: p < 0.0001). All statistical analyses were performed using GraphPad Prism software (Version 10) or ggpubr package in R.

## Acknowledgements

We thank Martine Nijs, Nena Dupont, and Kevin Feyen for technical assistance. We gratefully acknowledge the VIB flow core, the Olink service platform, and the lab of Diether Lambrechts for their help with RNA sequencing. We thank Wim Martinet for providing us with LC3B GFP-RFP mice. We also thank Esther Hoste for her help setting up the psoriasis model. This work was funded by the iBOF/21/053 Atlantis consortium for P.A. and G.B, the EOS MetaNiche consortium N° 40007532 for P.A., C.S., S.L., and G.B. G.B. also received support from the Flemish government FWO (G0A1122N). K.A.J. received a post-doctoral fellowship from the Flemish government FWO (12Y4322N). A.T. received support from an ERC starting grant (METATARGET, 101078355), an Emmy Noether Award from the German Research Foundation (DFG, 467788900), and the Ministry of Culture and Science of the State of North Rhine-Westphalia (NRW-Nachwuchsgruppenprogramm). W.S. received funds from the Deutsche Forschungsgemeinschaft (DFG, German Research Foundation)—Project-ID 422744262—TRR 289.

## Author contributions

K.A.J. and G.B. conceptualized the study. P.A.’s knowledge of autophagy and UPR provided essential input. K.A.J, P.A., and G.B. designed experiments, K.A.J., A.P.N., G.V., A.F.D.L, J.B., M.G., L.M., F.C., and G.A. performed experiments. C.S. and A.P.N. guided the intravital imaging experiments and provided valuable input, L.M. assisted in experimental setup. Y.H. conducted the murine scRNAseq analyses under supervision of F.R. and S.G. analyzed the human tonsil scRNAseq data. S.L. and N.V.B. provided human tonsillar tissue. W.S. and A.T. provided human psoriasis tissue. K.A.J. and G.B. wrote the manuscript; P.A. and A.T. edited the manuscript.

## Declaration of Interests

The authors declare no conflict of interest.

## Supplemental figure legends

**Figure S1:**
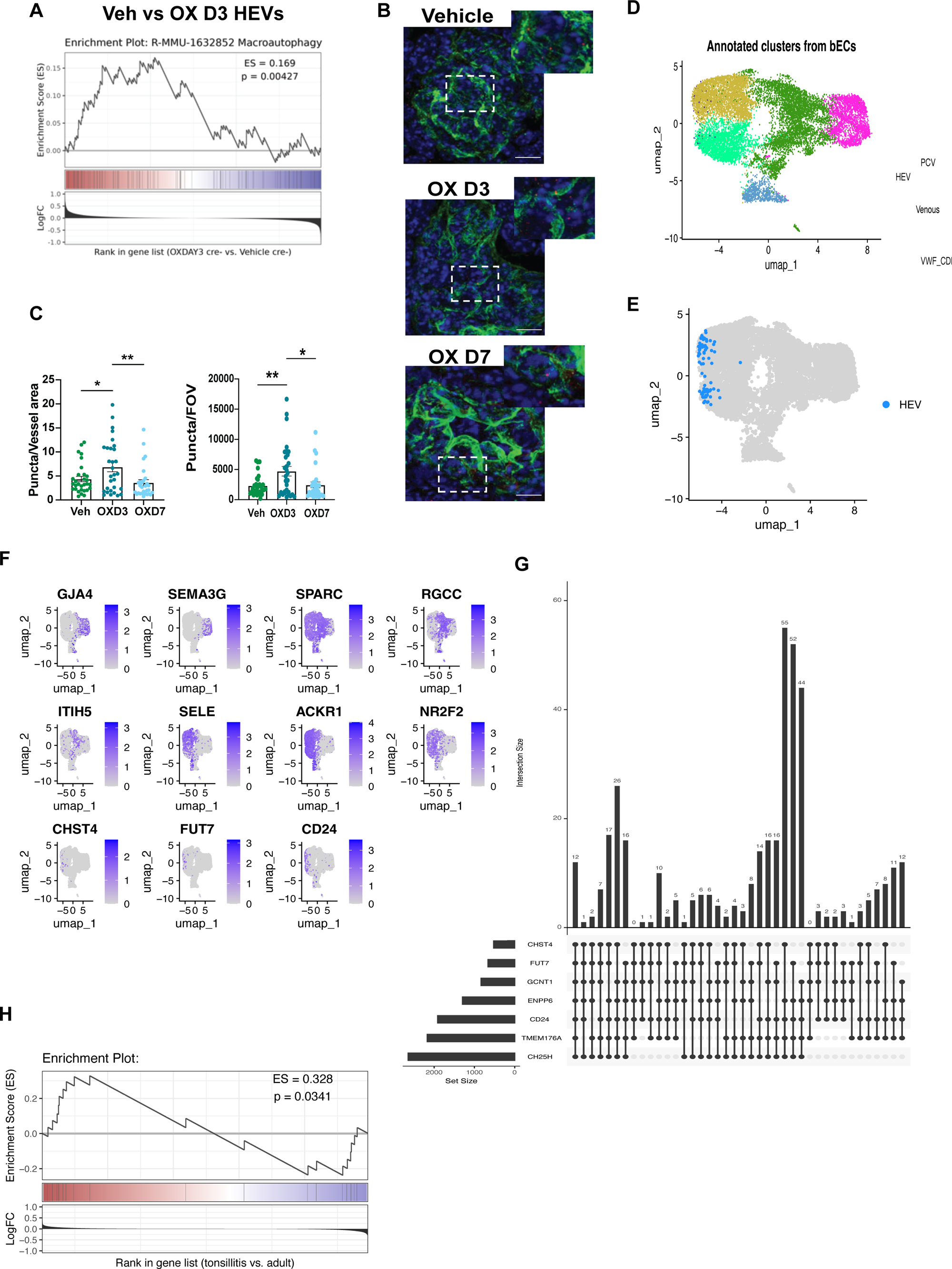
Autophagy is essential for HEVs during inflammation. (A) GSEA plot for macroautophagy in inflamed vs vehicle HEVs (B) Immunofluorescence staining of LC3B in vehicle, inflamed (OX D3) or recovering (OX D7) HEVs (Meca-79). Scale bar=10um (C) Quantifications of (B) (D) UMAP of EC clusters in Human tonsil samples (E) HEVs identified using HEV signature highlighted within UMAP (F) UMAPs showing top genes used to subcluster human tonsillar ECs (G) Upset plot showing the different gene combinations expressed in cells positive for HEV signature. (H) GSEA plot of active autophagy signature in Tonsilitis HEVs vs homeostatic HEVs. *p < 0.05; **p < 0.01; ***p < 0.001; ****p < 0.0001.

**Figure S2:**
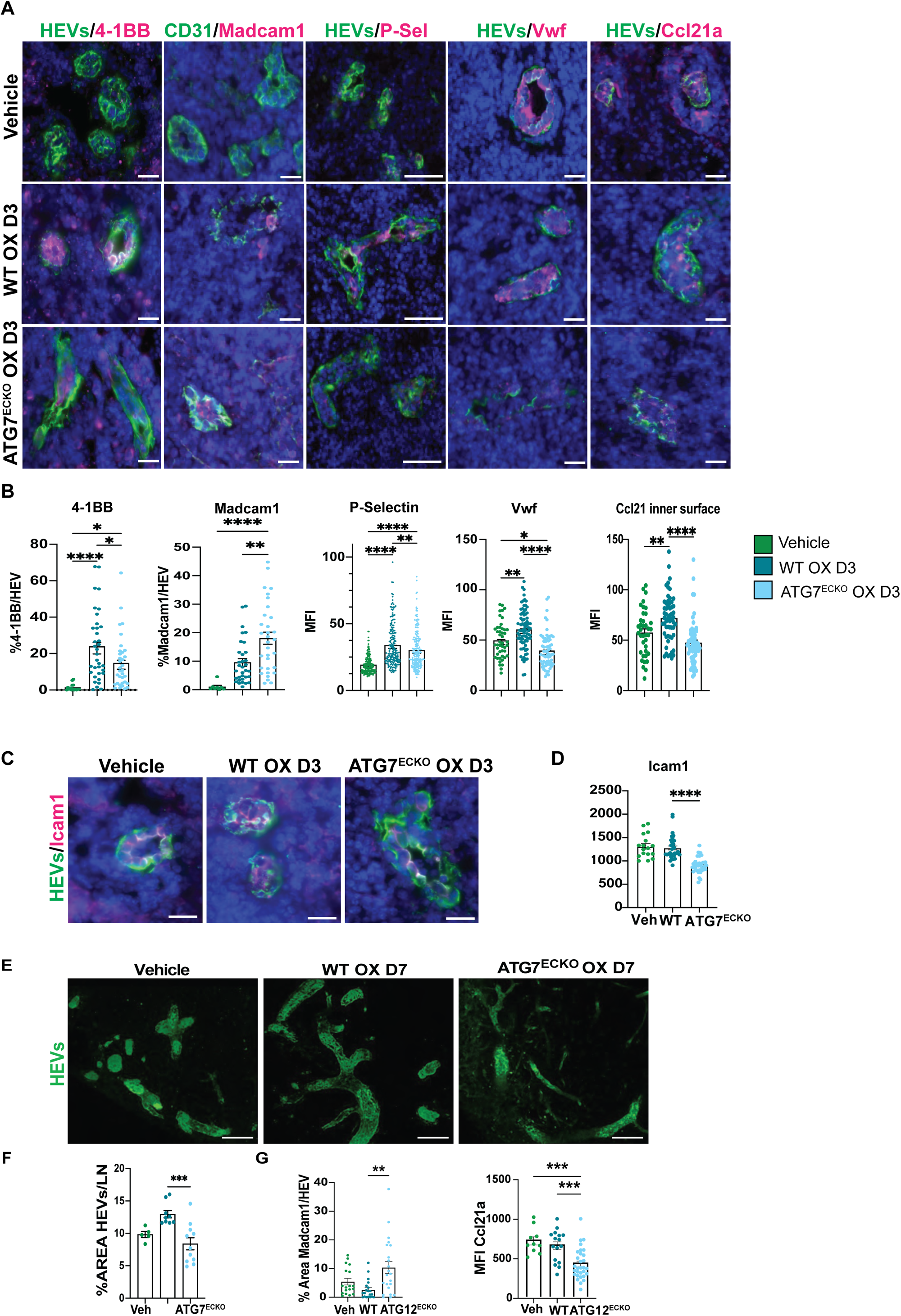
ATG7^ECKO^ Mice recapitulate ATG12^ECKO^ OX induced phenotype. (A) Immunofluorescence staining of HEVs, Cd31, P-selectin, 4-1BB, Madcam1, Ccl21a, Vwf in Vehicle or OX treated WT and ATG12^ECKO^ mice Scale bar for all but P-Selectin =20um, P-Selectin scale bar=50um (B) Quantification of stainings in (A) (C) Immunofluorescence staining of HEVs and Icam1 on D3 post OX treatment. Scale bar=20um (D) MFI quantification of (C) (E) 3D-reconstruction of immunofluorescence staining of HEVs on D7 post OX treatment. Scale bar=100um (F) Quantification of HEV area (Meca-79) per LN in (E) (G) Quantification of MFI of Ccl21 and percentage of Madcam1-positive HEVs in WT vs ATG12^ECKO^ mice on OX D7. *p < 0.05; **p < 0.01; ***p < 0.001; ****p < 0.0001.

**Figure S3:**
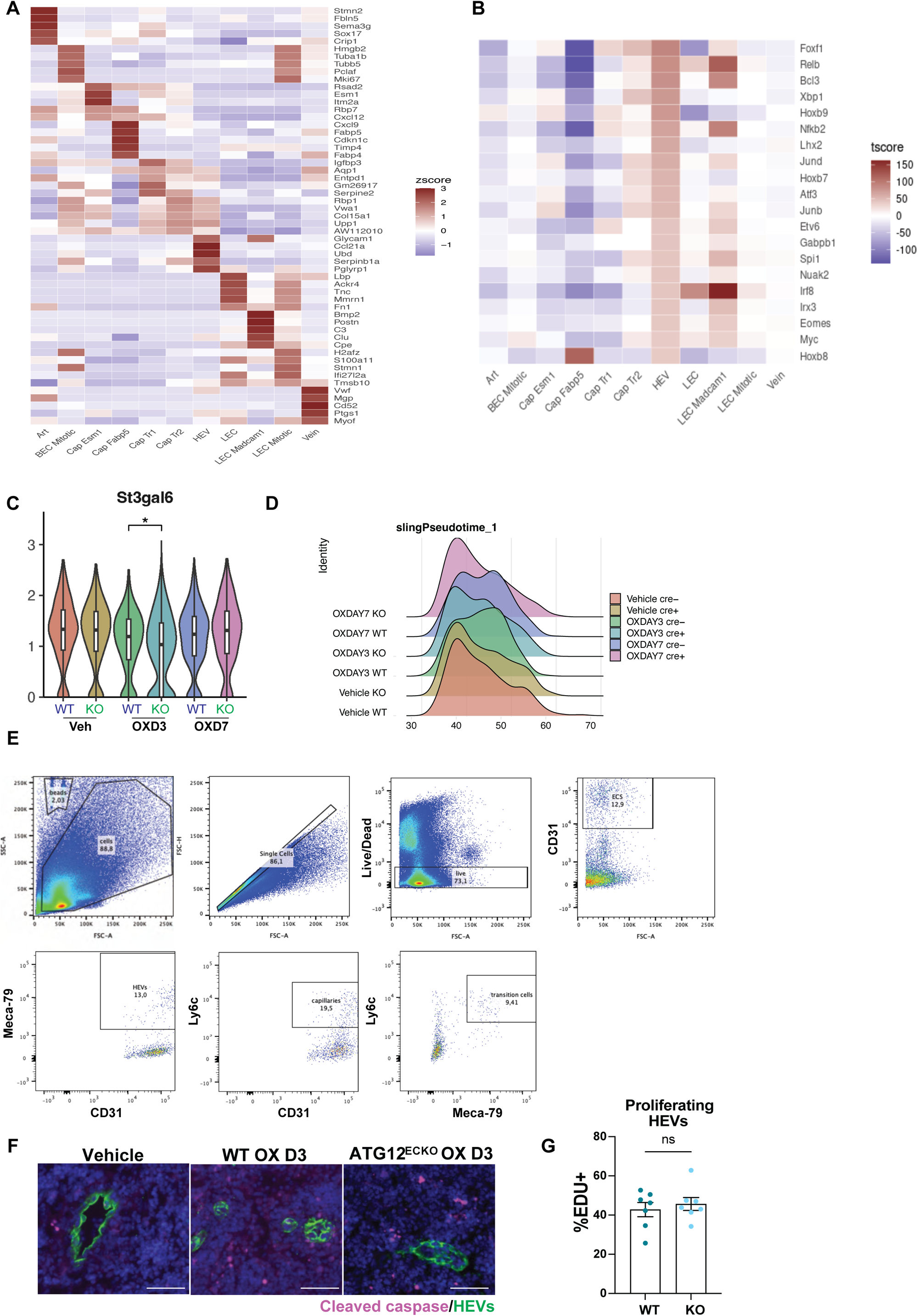
Loss of Autophagy in HEVs leads to dedifferentiation of these vessels. (A) Heatmap of top genes used for unsupervised clustering of ECs (B) Heatmap of top TFs in HEVs identified by SCENIC and their expression across EC clusters C) Violin Plots of selected enzyme involved in PNAd production (St3gal6) across samples (D) Graphical representation of Singshot analysis of HEV cluster plotted in pseudotime for WT and ATG12^ECKO^ on Vehicle, OX D3 and OX D7 (E) Gating of FACs analysis for double-positive HEVs (M79+ Ly6C+) in Figure 3H (F) Immunofluorescence staining of cleaved caspase in HEVs in Vehicle and OX D3 WT and ATG12^ECKO^ mice, Scale bar=50um (G) % of HEVs (Meca-79+) cells analyzed by FACs which are double positive for EDU on OX D3, each dot represents one mouse. *p < 0.05; **p < 0.01; ***p < 0.001; ****p < 0.0001.

**Figure S4:**
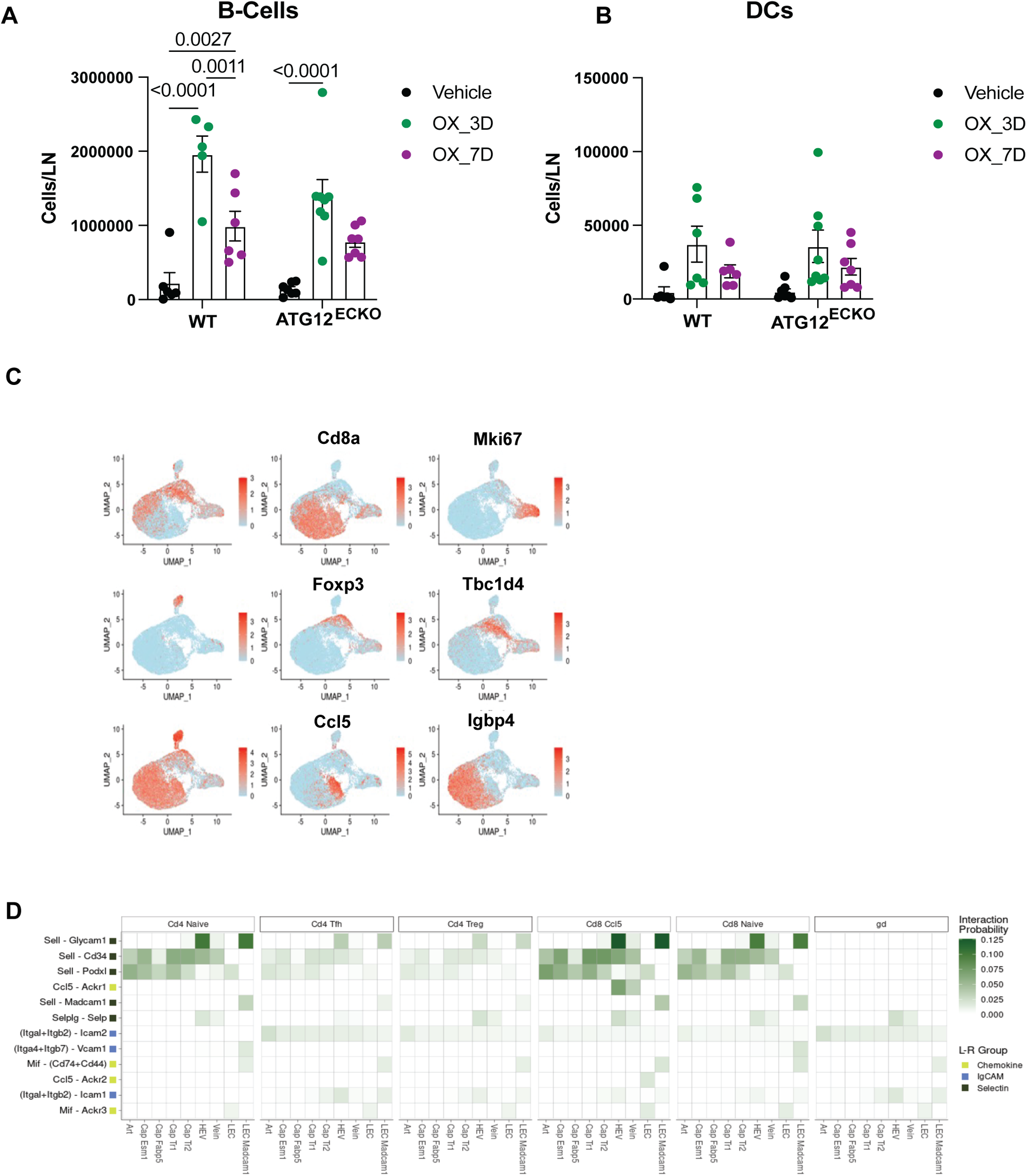
HEVs lose functionality in the absence of autophagy. (A-B) Characterization of B-cells and DCs in mouse LNs in WT and ATG12^ECKO^ samples treated with vehicle, OX D3 and OX D7 by FACs. Each dot represents one mouse. (C) UMAPs showing top genes used to subcluster T-Cells (D) Expanded predicted ligands (T-cells) and receptors (ECs) by CellChat

**Figure S5:**
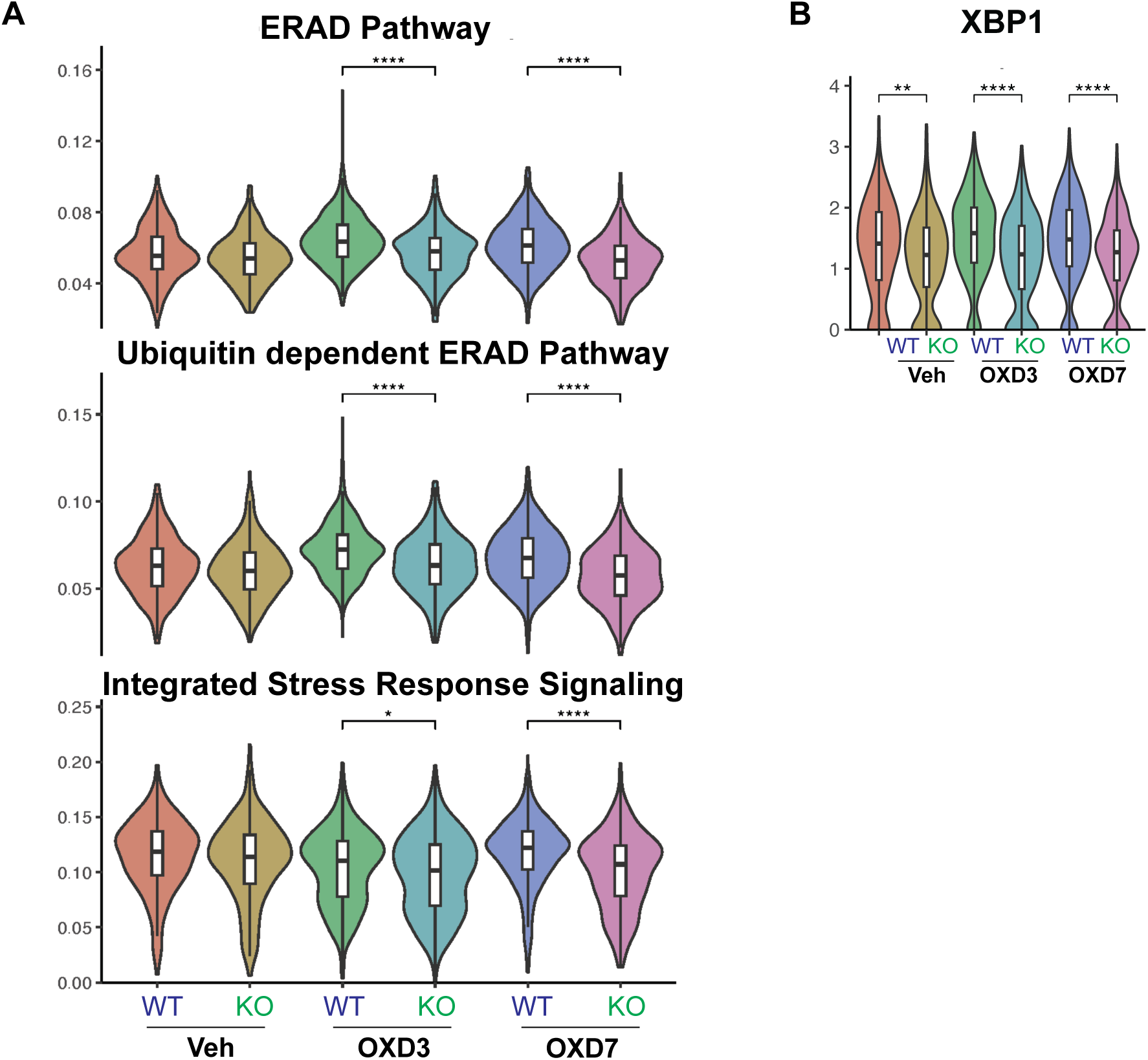
Autophagy affects ER related stress responses in HEVs. (A) Violin plots of signatures of ERAD, Ubiquitin-dependent ERAD, and the integrated stress response in HEVs across samples (B) Violin plot of Xbp1 expression in HEVs across samples.

**Figure S6:**
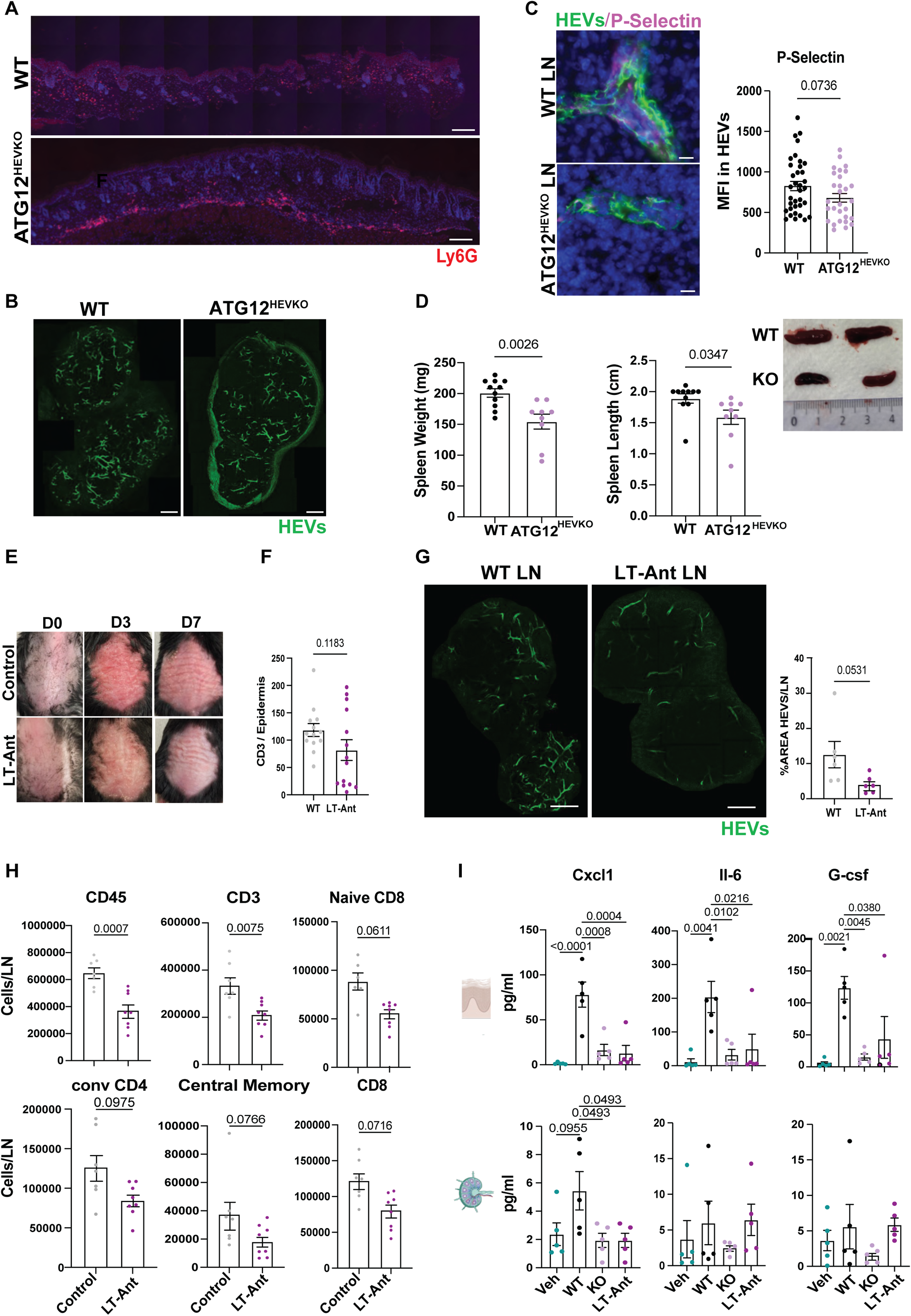
Blocking HEVs alleviates psoriasis phenotype. (A) Immunofluorescence staining of Ly6G (Neutrophils) in skin of WT and ATG12^HEVKO^ mice in psoriasis model. Scale bar= 250 um (B) Immunofluorescence staining of HEV area per LN in WT and ATG12^HEVKO^ mice in psoriasis model Scale Bar=250um (C) Quantification and images of spleen weight and length in WT and ATG12^HEVKO^ mice in psoriasis model. (D) Immunofluorescence staining and quantification of Mean fluorescence intensity of HEVs and P-Selectin in LNs of WT and ATG12^HEVKO^ mice in psoriasis model. Scale bar= 20um. (E) Images of psoriasis lesion in control vs. LTBR-antagonist (LT-Ant) treated mice on D0, D3, and D7 (F) Quantification of lymphocyte numbers per epidermis section (G) Immunofluorescence staining of HEVs and quantification of HEV area per LN in control and LT-Ant treated mice in psoriasis model. Scale bar =250 um (H) Immune cell number characterization of LNs from control and LT-Ant treated mice by FACs. Each dot represents one mouse. (I) Quantification of cytokine amount (pg/ml) in skin samples (top) and LN samples (bottom) for Vehicle, WT, ATG12 ^HEVKO^, and LTΒR-antag treated mice

## Notes

### Competing Interest Statement

The authors have declared no competing interest.

## References

1. Vella, G., Hua, Y., and Bergers, G. (2023). High endothelial venules in cancer: Regulation, function, and therapeutic implication. Cancer Cell 41, 527–545. 10.1016/j.ccell.2023.02.002.

2. Vella, G., Guelfi, S., and Bergers, G. (2021). High Endothelial Venules: A Vascular Perspective on Tertiary Lymphoid Structures in Cancer. Front. Immunol. 12. 10.3389/fimmu.2021.736670.

3. Blanchard, L., and Girard, J.-P. (2021). High endothelial venules (HEVs) in immunity, inflammation and cancer. Angiogenesis 24, 719–753. 10.1007/s10456-021-09792-8.

4. Veerman, K., Tardiveau, C., Martins, F., Coudert, J., and Girard, J.-P. (2019). Single-Cell Analysis Reveals Heterogeneity of High Endothelial Venules and Different Regulation of Genes Controlling Lymphocyte Entry to Lymph Nodes. Cell Rep 26, 3116–3131.e5. 10.1016/j.celrep.2019.02.042.

5. Xu, B., Wagner, N., Pham, L.N., Magno, V., Shan, Z., Butcher, E.C., and Michie, S.A. (2003). Lymphocyte homing to bronchus-associated lymphoid tissue (BALT) is mediated by L-selectin/PNAd, alpha4beta1 integrin/VCAM-1, and LFA-1 adhesion pathways. J Exp Med 197, 1255–1267. 10.1084/jem.20010685.

6. Thaunat, O., Patey, N., Caligiuri, G., Gautreau, C., Mamani-Matsuda, M., Mekki, Y., Dieu-Nosjean, M.-C., Eberl, G., Ecochard, R., Michel, J.-B., et al. (2010). Chronic rejection triggers the development of an aggressive intragraft immune response through recapitulation of lymphoid organogenesis. J Immunol 185, 717–728. 10.4049/jimmunol.0903589.

7. Kobayashi, M., Mitoma, J., Nakamura, N., Katsuyama, T., Nakayama, J., and Fukuda, M. (2004). Induction of peripheral lymph node addressin in human gastric mucosa infected by Helicobacter pylori. Proceedings of the National Academy of Sciences 101, 17807–17812. 10.1073/pnas.0407503101.

8. Hua, Y., Vella, G., Rambow, F., Allen, E., Antoranz Martinez, A., Duhamel, M., Takeda, A., Jalkanen, S., Junius, S., Smeets, A., et al. (2022). Cancer immunotherapies transition endothelial cells into HEVs that generate TCF1+ T lymphocyte niches through a feed-forward loop. Cancer Cell 40, 1600–1618.e10. 10.1016/j.ccell.2022.11.002.

9. Helmink, B.A., Reddy, S.M., Gao, J., Zhang, S., Basar, R., Thakur, R., Yizhak, K., Sade-Feldman, M., Blando, J., Han, G., et al. (2020). B cells and tertiary lymphoid structures promote immunotherapy response. Nature 577, 549–555. 10.1038/s41586-019-1922-8.

10. Budair, F.M., Nomura, T., Hirata, M., and Kabashima, K. (2024). PNAd-expressing vessels characterize the dermis of CD3+ T-cell-mediated cutaneous diseases. Clin Exp Immunol 216, 80–88. 10.1093/cei/uxae003.

11. Jeucken, K.C.M., Koning, J.J., Mebius, R.E., and Tas, S.W. (2019). The Role of Endothelial Cells and TNF-Receptor Superfamily Members in Lymphoid Organogenesis and Function During Health and Inflammation. Front. Immunol. 10. 10.3389/fimmu.2019.02700.

12. Dikic, I., and Elazar, Z. (2018). Mechanism and medical implications of mammalian autophagy. Nat Rev Mol Cell Biol 19, 349–364. 10.1038/s41580-018-0003-4.

13. Deretic, V. (2021). Autophagy in inflammation, infection, and immunometabolism. Immunity 54, 437–453. 10.1016/j.immuni.2021.01.018.

14. Reglero-Real, N., Pérez-Gutiérrez, L., Yoshimura, A., Rolas, L., Garrido-Mesa, J., Barkaway, A., Pickworth, C., Saleeb, R.S., Gonzalez-Nuñez, M., Austin-Williams, S.N., et al. (2021). Autophagy modulates endothelial junctions to restrain neutrophil diapedesis during inflammation. Immunity 54, 1989–2004.e9. 10.1016/j.immuni.2021.07.012.

15. Verhoeven, J., Jacobs, K.A., Rizzollo, F., Lodi, F., Hua, Y., Poźniak, J., Narayanan Srinivasan, A., Houbaert, D., Shankar, G., More, S., et al. (2023). Tumor endothelial cell autophagy is a key vascular-immune checkpoint in melanoma. EMBO Molecular Medicine 15, e18028. 10.15252/emmm.202318028.

16. Kwon, J., Kim,Jihyun, and and Kim, K.I. (2023). Crosstalk between endoplasmic reticulum stress response and autophagy in human diseases. Animal Cells and Systems 27, 29–37. 10.1080/19768354.2023.2181217.

17. B’chir, W., Maurin, A.-C., Carraro, V., Averous, J., Jousse, C., Muranishi, Y., Parry, L., Stepien, G., Fafournoux, P., and Bruhat, A. (2013). The eIF2α/ATF4 pathway is essential for stress-induced autophagy gene expression. Nucleic Acids Res 41, 7683–7699. 10.1093/nar/gkt563.

18. Vidal, R.L., Figueroa, A., Court, F.A., Thielen, P., Molina, C., Wirth, C., Caballero, B., Kiffin, R., Segura-Aguilar, J., Cuervo, A.M., et al. (2012). Targeting the UPR transcription factor XBP1 protects against Huntington’s disease through the regulation of FoxO1 and autophagy. Hum Mol Genet 21, 2245–2262. 10.1093/hmg/dds040.

19. Nishimura, T., and Tooze, S.A. (2020). Emerging roles of ATG proteins and membrane lipids in autophagosome formation. Cell Discov 6, 32. 10.1038/s41421-020-0161-3.

20. Liao, S., and Ruddle, N.H. (2006). Synchrony of High Endothelial Venules and Lymphatic Vessels Revealed by Immunization1. The Journal of Immunology 177, 3369–3379. 10.4049/jimmunol.177.5.3369.

21. Lin, F., Wang, Z.V., and Hill, J.A. (2014). Seeing is believing. Autophagy 10, 691–693. 10.4161/auto.27749.

22. Li, L., Wang, Z.V., Hill, J.A., and Lin, F. (2014). New Autophagy Reporter Mice Reveal Dynamics of Proximal Tubular Autophagy. J Am Soc Nephrol 25, 305–315. 10.1681/ASN.2013040374.

23. Bordi, M., De Cegli, R., Testa, B., Nixon, R.A., Ballabio, A., and Cecconi, F. (2021). A gene toolbox for monitoring autophagy transcription. Cell Death Dis 12, 1–7. 10.1038/s41419-021-04121-9.

24. Mebius, R.E., Streeter, P.R., Michie, S., Butcher, E.C., and Weissman, I.L. (1996). A developmental switch in lymphocyte homing receptor and endothelial vascular addressin expression regulates lymphocyte homing and permits CD4+ CD3-cells to colonize lymph nodes. Proceedings of the National Academy of Sciences 93, 11019– 11024. 10.1073/pnas.93.20.11019.

25. Verhoeven, J., Agostinis, P., and Agrawal, M. (2023). Methods for Isolation of Tumor-Associated Endothelial Cells for Surface Protein Analysis and Sorting by Flowcytometry. Methods Mol Biol 2572, 45–54. 10.1007/978-1-0716-2703-7_3.

26. Brulois, K., Rajaraman, A., Szade, A., Nordling, S., Bogoslowski, A., Dermadi, D., Rahman, M., Kiefel, H., O’Hara, E., Koning, J.J., et al. (2020). A molecular map of murine lymph node blood vascular endothelium at single cell resolution. Nat Commun 11, 3798. 10.1038/s41467-020-17291-5.

27. Browning, J.L., Allaire, N., Ngam-Ek, A., Notidis, E., Hunt, J., Perrin, S., and Fava, R.A. (2005). Lymphotoxin-beta receptor signaling is required for the homeostatic control of HEV differentiation and function. Immunity 23, 539–550. 10.1016/j.immuni.2005.10.002.

28. Senft, D., and Ronai, Z.A. (2015). UPR, autophagy, and mitochondria crosstalk underlies the ER stress response. Trends in Biochemical Sciences 40, 141–148. 10.1016/j.tibs.2015.01.002.

29. Marchetti, L., Francisco, D., Soldati, S., Haghayegh Jahromi, N., Barcos, S., Gruber, I., Pareja, J.R., Thiriot, A., von Andrian, U., Deutsch, U., et al. (2022). ACKR1 favors transcellular over paracellular T-cell diapedesis across the blood-brain barrier in neuroinflammation in vitro. Eur J Immunol 52, 161–177. 10.1002/eji.202149238.

30. Girbl, T., Lenn, T., Perez, L., Rolas, L., Barkaway, A., Thiriot, A., Fresno, C. del, Lynam, E., Hub, E., Thelen, M., et al. (2018). Distinct Compartmentalization of the Chemokines CXCL1 and CXCL2 and the Atypical Receptor ACKR1 Determine Discrete Stages of Neutrophil Diapedesis. Immunity 49, 1062–1076.e6. 10.1016/j.immuni.2018.09.018.

31. Moussion, C., and Girard, J.-P. (2011). Dendritic cells control lymphocyte entry to lymph nodes through high endothelial venules. Nature 479, 542–546. 10.1038/nature10540.

32. van de Pavert, S.A., and Mebius, R.E. (2010). New insights into the development of lymphoid tissues. Nat Rev Immunol 10, 664–674. 10.1038/nri2832.

33. Peske, J.D., Thompson, E.D., Gemta, L., Baylis, R.A., Fu, Y.-X., and Engelhard, V.H. (2015). Effector lymphocyte-induced lymph node-like vasculature enables naive T-cell entry into tumours and enhanced anti-tumour immunity. Nat Commun 6, 7114. 10.1038/ncomms8114.

34. Sun, S.-C. (2017). The non-canonical NF-κB pathway in immunity and inflammation. Nat Rev Immunol 17, 545–558. 10.1038/nri.2017.52.

35. Newman, A.C., Kemp, A.J., Drabsch, Y., Behrends, C., and Wilkinson, S. (2017). Autophagy acts through TRAF3 and RELB to regulate gene expression via antagonism of SMAD proteins. Nat Commun 8, 1537. 10.1038/s41467-017-00859-z.

36. Leng, H., Simon, A.K., and Horwood, N.J. (2024). Blocking glycosphingolipid production alters autophagy in osteoclasts and improves myeloma bone disease. Autophagy 20, 930–932. 10.1080/15548627.2023.2208931.

37. Norton, P., Comunale, M.A., Herrera, H., Wang, M., Romano, P.R., and Mehta, A. (2016). Development and application of a novel recombinant Aleuria aurantia lectin with enhanced core fucose binding for identification of glycoprotein biomarker of hepatocellular carcinoma. Proteomics 16, 3126–3136. 10.1002/pmic.201600064.

38. Hetz, C., Zhang, K., and Kaufman, R.J. (2020). Mechanisms, regulation and functions of the unfolded protein response. Nat Rev Mol Cell Biol 21, 421–438. 10.1038/s41580-020-0250-z.

39. Bi, Y., Brulois, K., Ayesha, A., Xiang, M., Ballet, R., Ocón, B., Dinh, T., Lazarus, N., Kunte, M., Wang, Y., et al. (2025). Common Gene Networks Orchestrate Organelle Architecture and Inter-Organelle Metabolic Flows for Mucin Production in High Endothelial and Goblet Cells. bioRxiv, 2025.06.19.660616. 10.1101/2025.06.19.660616.

40. Jabeen, M., Boisgard, A.-S., Danoy, A., El Kholti, N., Salvi, J.-P., Boulieu, R., Fromy, B., Verrier, B., and Lamrayah, M. (2020). Advanced Characterization of Imiquimod-Induced Psoriasis-Like Mouse Model. Pharmaceutics 12, 789. 10.3390/pharmaceutics12090789.

41. Houbaert, D., Nikolakopoulos, A.P., Jacobs, K.A., Meçe, O., Roels, J., Shankar, G., Agrawal, M., More, S., Ganne, M., Rillaerts, K., et al. (2024). An autophagy program that promotes T cell egress from the lymph node controls responses to immune checkpoint blockade. Cell Reports 43, 114020. 10.1016/j.celrep.2024.114020.

42. Zhou, X., Chen, Y., Cui, L., Shi, Y., and Guo, C. (2022). Advances in the pathogenesis of psoriasis: from keratinocyte perspective. Cell Death Dis 13, 1–13. 10.1038/s41419-022-04523-3.

43. Brembilla, N.C., and Boehncke, W.-H. (2023). Revisiting the interleukin 17 family of cytokines in psoriasis: pathogenesis and potential targets for innovative therapies. Front. Immunol. 14. 10.3389/fimmu.2023.1186455.

44. Wolk, K., Witte, E., Wallace, E., Döcke, W.-D., Kunz, S., Asadullah, K., Volk, H.-D., Sterry, W., and Sabat, R. (2006). IL-22 regulates the expression of genes responsible for antimicrobial defense, cellular differentiation, and mobility in keratinocytes: a potential role in psoriasis. Eur J Immunol 36, 1309–1323. 10.1002/eji.200535503.

45. Onder, L., Danuser, R., Scandella, E., Firner, S., Chai, Q., Hehlgans, T., Stein, J.V., and Ludewig, B. (2013). Endothelial cell-specific lymphotoxin-β receptor signaling is critical for lymph node and high endothelial venule formation. J Exp Med 210, 465–473. 10.1084/jem.20121462.

46. Seino, J., Wang, L., Harada, Y., Huang, C., Ishii, K., Mizushima, N., and Suzuki, T. (2013). Basal Autophagy Is Required for the Efficient Catabolism of Sialyloligosaccharides *. Journal of Biological Chemistry 288, 26898–26907. 10.1074/jbc.M113.464503.

47. Guo, B., Liang, Q., Li, L., Hu, Z., Wu, F., Zhang, P., Ma, Y., Zhao, B., Kovács, A.L., Zhang, Z., et al. (2014). O-GlcNAc-modification of SNAP-29 regulates autophagosome maturation. Nat Cell Biol 16, 1215–1226. 10.1038/ncb3066.

48. Pyo, K.E., Kim, C.R., Lee, M., Kim, J.-S., Kim, K.I., and Baek, S.H. (2018). ULK1 O-GlcNAcylation Is Crucial for Activating VPS34 via ATG14L during Autophagy Initiation. Cell Reports 25, 2878–2890.e4. 10.1016/j.celrep.2018.11.042.

49. B’chir, W., Maurin, A.-C., Carraro, V., Averous, J., Jousse, C., Muranishi, Y., Parry, L., Stepien, G., Fafournoux, P., and Bruhat, A. (2013). The eIF2α/ATF4 pathway is essential for stress-induced autophagy gene expression. Nucleic Acids Research 41, 7683–7699. 10.1093/nar/gkt563.

50. Stress-induced self-cannibalism: on the regulation of autophagy by endoplasmic reticulum stress | Cellular and Molecular Life Sciences https://link.springer.com/article/10.1007/s00018-012-1173-4?utm_source=getftr&utm_medium=getftr&utm_campaign=getftr_pilot&getft_integrator=sciencedirect_contenthosting.

51. Quan, W., Hur, K.Y., Lim, Y., Oh, S.H., Lee, J.-C., Kim, K.H., Kim, G.H., Kim, S.-W., Kim, H.L., Lee, M.-K., et al. (2012). Autophagy deficiency in beta cells leads to compromised unfolded protein response and progression from obesity to diabetes in mice. Diabetologia 55, 392–403. 10.1007/s00125-011-2350-y.

52. Xu, J., Zhu, R., Chen, Y., Li, X., Chen, X., Liu, M., and Huang, Y. (2024). Chloroquine-induced proinsulin misfolding in the endoplasmic reticulum underlies the attenuation of mature insulin synthesis. The FASEB Journal 38, e70201. 10.1096/fj.202401945R.

53. Quan, W., Hur, K.Y., Lim, Y., Oh, S.H., Lee, J.-C., Kim, K.H., Kim, G.H., Kim, S.-W., Kim, H.L., Lee, M.-K., et al. (2012). Autophagy deficiency in beta cells leads to compromised unfolded protein response and progression from obesity to diabetes in mice. Diabetologia 55, 392–403. 10.1007/s00125-011-2350-y.

54. Shaffer, A.L., Shapiro-Shelef, M., Iwakoshi, N.N., Lee, A.-H., Qian, S.-B., Zhao, H., Yu, X., Yang, L., Tan, B.K., Rosenwald, A., et al. (2004). XBP1, downstream of Blimp-1, expands the secretory apparatus and other organelles, and increases protein synthesis in plasma cell differentiation. Immunity 21, 81–93. 10.1016/j.immuni.2004.06.010.

55. Yang, M., Luo, S., Wang, X., Li, C., Yang, J., Zhu, X., Xiao, L., and Sun, L. (2021). ER-Phagy: A New Regulator of ER Homeostasis. Front. Cell Dev. Biol. 9. 10.3389/fcell.2021.684526.

56. Armstrong, A.W., Mehta, M.D., Schupp, C.W., Gondo, G.C., Bell, S.J., and Griffiths, C.E.M. (2021). Psoriasis Prevalence in Adults in the United States. JAMA Dermatology 157, 940– 946. 10.1001/jamadermatol.2021.2007.

57. Lee, H.-J., and Kim, M. (2023). Challenges and Future Trends in the Treatment of Psoriasis. Int J Mol Sci 24, 13313. 10.3390/ijms241713313.

58. Kogame, T., Yamashita, R., Hirata, M., Kataoka, T.R., Kamido, H., Ueshima, C., Matsui, M., Nomura, T., and Kabashima, K. (2018). Analysis of possible structures of inducible skin-associated lymphoid tissue in lupus erythematosus profundus. J Dermatol 45, 1117–1121. 10.1111/1346-8138.14498.

59. Malhotra, R., Warne, J.P., Salas, E., Xu, A.W., and Debnath, J. (2015). Loss of Atg12, but not Atg5, in pro-opiomelanocortin neurons exacerbates diet-induced obesity. Autophagy.

60. Sörensen, I., Adams, R.H., and Gossler, A. (2009). DLL1-mediated Notch activation regulates endothelial identity in mouse fetal arteries. Blood 113, 5680–5688. 10.1182/blood-2008-08-174508.

61. Messal, H.A., van Rheenen, J., and Scheele, C.L.G.J. (2021). An Intravital Microscopy Toolbox to Study Mammary Gland Dynamics from Cellular Level to Organ Scale. J Mammary Gland Biol Neoplasia 26, 9–27. 10.1007/s10911-021-09487-2.

62. Menzel, L., Zschummel, M., and Rehm, A. (2022). Analyses of murine lymph node endothelial cell subsets using single-cell RNA sequencing and spectral flow cytometry. STAR Protoc 3, 101267. 10.1016/j.xpro.2022.101267.

63. Hao, Y., Hao, S., Andersen-Nissen, E., Mauck, W.M., Zheng, S., Butler, A., Lee, M.J., Wilk, A.J., Darby, C., Zager, M., et al. (2021). Integrated analysis of multimodal single-cell data. Cell 184, 3573–3587.e29. 10.1016/j.cell.2021.04.048.

64. Korsunsky, I., Millard, N., Fan, J., Slowikowski, K., Zhang, F., Wei, K., Baglaenko, Y., Brenner, M., Loh, P.-R., and Raychaudhuri, S. (2019). Fast, sensitive and accurate integration of single-cell data with Harmony. Nat Methods 16, 1289–1296. 10.1038/s41592-019-0619-0.

65. Aibar, S., González-Blas, C.B., Moerman, T., Huynh-Thu, V.A., Imrichova, H., Hulselmans, G., Rambow, F., Marine, J.-C., Geurts, P., Aerts, J., et al. (2017). SCENIC: single-cell regulatory network inference and clustering. Nat Methods 14, 1083–1086. 10.1038/nmeth.4463.

66. Setty, M., Kiseliovas, V., Levine, J., Gayoso, A., Mazutis, L., and Pe’er, D. (2019). Characterization of cell fate probabilities in single-cell data with Palantir. Nat Biotechnol 37, 451–460. 10.1038/s41587-019-0068-4.

67. Bergen, V., Lange, M., Peidli, S., Wolf, F.A., and Theis, F.J. (2020). Generalizing RNA velocity to transient cell states through dynamical modeling. Nat Biotechnol 38, 1408– 1414. 10.1038/s41587-020-0591-3.

68. Gulati, G.S., Sikandar, S.S., Wesche, D.J., Manjunath, A., Bharadwaj, A., Berger, M.J., Ilagan, F., Kuo, A.H., Hsieh, R.W., Cai, S., et al. (2020). Single-cell transcriptional diversity is a hallmark of developmental potential. Science 367, 405–411. 10.1126/science.aax0249.

69. Hua, Y., Weng, L., Zhao, F., and Rambow, F. (2024). SeuratExtend: Streamlining Single-Cell RNA-Seq Analysis Through an Integrated and Intuitive Framework. Preprint at bioRxiv, 10.1101/2024.08.01.606144 10.1101/2024.08.01.606144.

70. Jin, S., Plikus, M.V., and Nie, Q. (2025). CellChat for systematic analysis of cell-cell communication from single-cell transcriptomics. Nat Protoc 20, 180–219. 10.1038/s41596-024-01045-4.

71. De Martin, A., Stanossek, Y., Lütge, M., Cadosch, N., Onder, L., Cheng, H.-W., Brandstadter, J.D., Maillard, I., Stoeckli, S.J., Pikor, N.B., et al. (2023). PI16+ reticular cells in human palatine tonsils govern T cell activity in distinct subepithelial niches. Nat Immunol 24, 1138–1148. 10.1038/s41590-023-01502-4.

72. Hao, Y., Stuart, T., Kowalski, M.H., Choudhary, S., Hoffman, P., Hartman, A., Srivastava, A., Molla, G., Madad, S., Fernandez-Granda, C., et al. (2024). Dictionary learning for integrative, multimodal and scalable single-cell analysis. Nat Biotechnol 42, 293–304. 10.1038/s41587-023-01767-y.

73. Li, Q., Zhu, Z., Wang, L., Lin, Y., Fang, H., Lei, J., Cao, T., Wang, G., and Dang, E. (2021). Single-cell transcriptome profiling reveals vascular endothelial cell heterogeneity in human skin. Theranostics 11, 6461–6476. 10.7150/thno.54917.

74. Jassal, B., Matthews, L., Viteri, G., Gong, C., Lorente, P., Fabregat, A., Sidiropoulos, K., Cook, J., Gillespie, M., Haw, R., et al. (2020). The reactome pathway knowledgebase. Nucleic Acids Res 48, D498–D503. 10.1093/nar/gkz1031.

75. L. Lun, A.T., Bach, K., and Marioni, J.C. (2016). Pooling across cells to normalize single-cell RNA sequencing data with many zero counts. Genome Biology 17, 75. 10.1186/s13059-016-0947-7.

